# The 3D structure of lipidic fibrils of α-synuclein

**DOI:** 10.1101/2022.03.02.481946

**Authors:** Benedikt Frieg, Leif Antonschmidt, Christian Dienemann, James A. Geraets, Eszter E. Najbauer, Dirk Matthes, Bert L. de Groot, Loren B. Andreas, Stefan Becker, Christian Griesinger, Gunnar F. Schröder

## Abstract

α-synuclein (αSyn) is abundant in neurons, but its misfolding and abnormal fibrillization are associated with severe neurodegenerative diseases. Although interactions between αSyn and phospholipid membranes are relevant during αSyn fibril assembly, insights into the interactions of αSyn fibrils with phospholipids have remained elusive. Here, we present six novel polymorphic atomic structures of αSyn fibrils aggregated in the presence of phospholipids. The structures reveal that phospholipids favor a novel protofilament fold, mediate an unusual arrangement of protofilaments, and fill the central cavities between the protofilaments. These findings provide a structural rationale for fibril-induced lipid extraction, a mechanism likely to be involved in the development of α-synucleinopathies.

## Main Text

The pathological causes for α-synucleinopathies, including Parkinson’s disease (PD), multiple system atrophy, and dementia with Lewy bodies (DLB), are largely unknown, but accumulating evidence suggests that these diseases are related to the misfolding and aggregation of α-synuclein (αSyn) into amyloid fibrils (*1-3*). A pathological hallmark for PD and DLB is the presence of large neuronal inclusions called Lewy bodies (LB) and αSyn fibrils have been identified as one constituent of LB (*4-6*). Several studies on the composition of LB extracted postmortem from brain tissue of PD-confirmed patients revealed that lipids and membranous organelles are a significant component of LB (*7-9*). Although molecular interactions between αSyn and lipids have been discussed as relevant for PD pathology for decades (*10*), detailed structural characterization is desirable. Here, we report cryo-electron microscopy (cryo-EM) structures of αSyn fibrils grown in the presence of phospholipid vesicles and used molecular dynamics (MD) simulations together with published Nuclear Magnetic Resonance (NMR) data confirming specific lipid-fibril interactions.

*De novo* aggregation in a 1:1 mixture of 1-palmitoyl-2-oleoyl-*sn*-glycero-3-phosphate (POPA) and 1-palmitoyl-2-oleoyl-*sn*-glycero-3-phosphocholine (POPC) with a 5:1 lipid to protein ratio was induced by sonication under protein misfolding cyclic amplification conditions and completed under gentle orbital shaking to elongate the fibrils (*11*) (**Fig. S1**). The lipids were selected as a simplification of negatively charged synaptic vesicles (*12*) to recapitulate the established binding of monomeric αSyn to lipids via its N-terminus (*13, 14*). We confirmed the presence of αSyn fibrils by cryo-EM screening and selected three preparations of αSyn fibrils for which independent image data sets were collected. Extensive classification during 3D reconstruction revealed three dominant protofilament folds (*L1, L2*, and *L3*) that form in total six different fibrils. Short sonication periods favor fibrils of the *L1* fold (**Fig. 1A**-**E**), while extensive sonication is needed to yield larger populations of *L2* and *L3* fibrils (**Fig. 2A-F, S1, S2**).

**Fig. 1.**
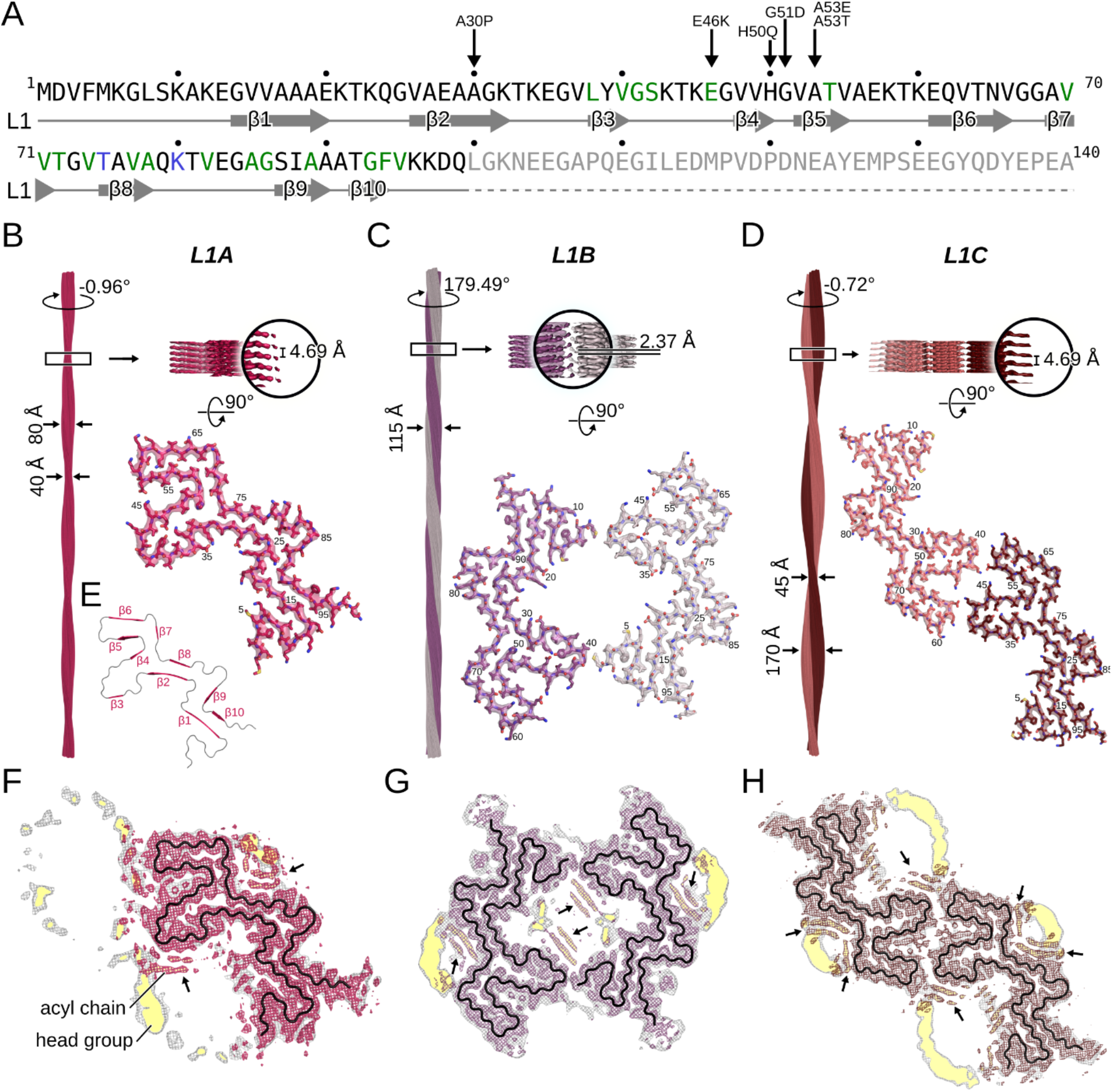
Cryo-EM structures of the *L1* αSyn fibrils. **A:** Sequence and secondary structure of human aSyn. Familial PD mutation sites (black arrow) localized within the lipid binding N-terminal region (residues 1–60) (*32*). Green-colored residues bind to the lipid acyl chain, blue to the choline moiety (*11*), and gray were not resolved. **B-D:** Cryo-EM structures of *L1A* (**B**), *L1B* (**C**), and *L1C* (**D**) fibrils (protofilaments colored differently). The atomic models are shown as sticks. Labels denote the fibril width, the helical twist and rise, and residue numbers. The density maps in the lower panels are displayed using the carve feature in PyMOL at a distance of 2 Å. **E**: Backbone of the *L1* fibrils with the β1 - β10 colored magenta and loops in gray. **F-H:** Overlay of a sharpened high-resolution map shown in magenta (**F**), purple (**G**), and brown (**H**) and an unsharpened, 4.5 Å low-pass filtered density in gray. The backbone is shown as a black ribbon. Densities highlighted with a yellow background are reminiscent of lipid micelles.

**Fig. 2.**
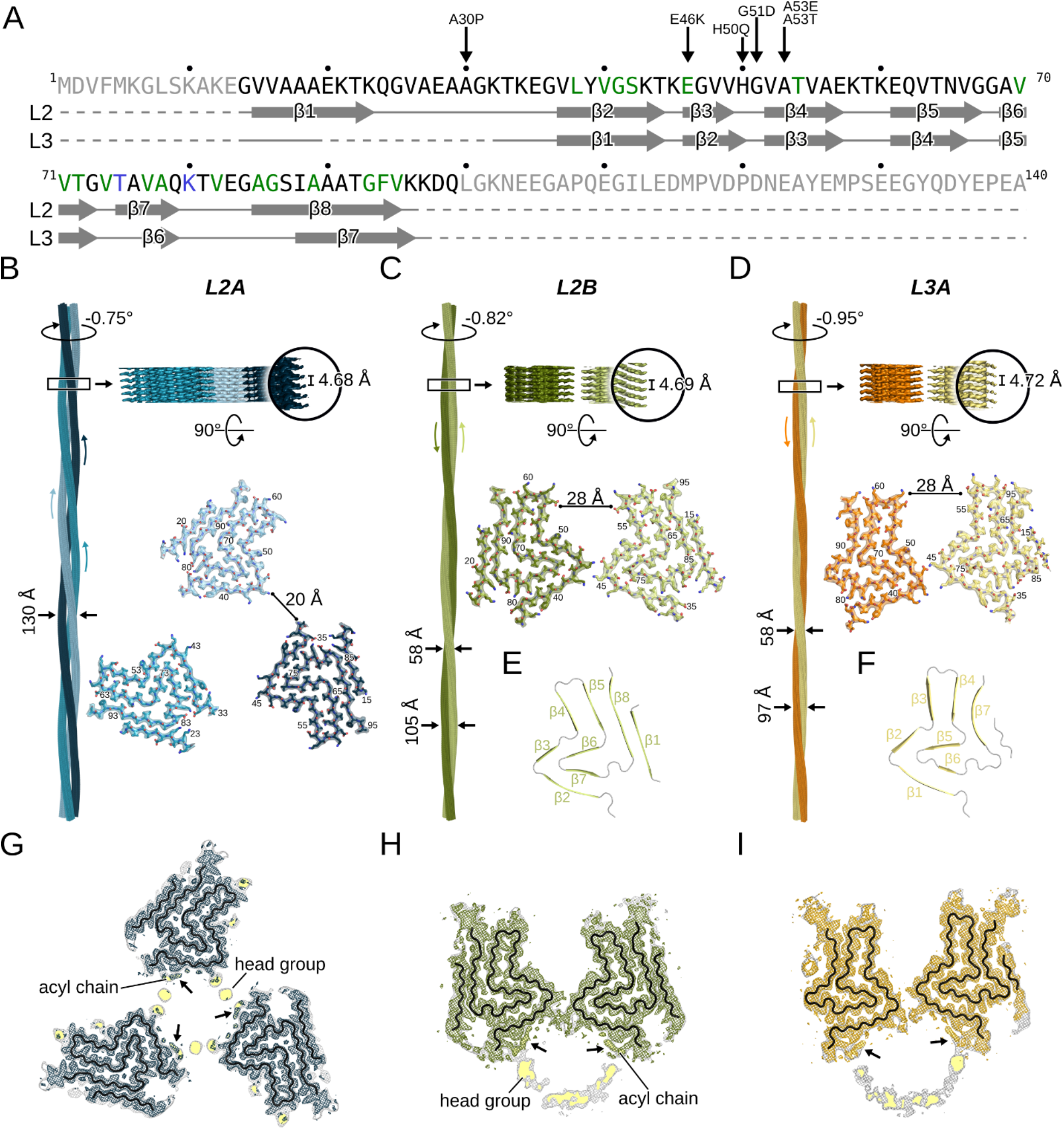
Cryo-EM structures of *L2* and *L3* αSyn fibrils. **A:** See **Fig. 1A** for details. **B-D:** Cryo-EM structures of *L2A* (**B**), *L2B* (**C**), and *L3A* (**D**) fibrils (protofilaments colored differently). The atomic models are shown as sticks. Labels denote the fibril width, the helical twist and rise, and residue numbers. The density maps in the lower panels are displayed using the carve feature in PyMOL at a distance of 2 Å. **E, F**: Backbone trace of the *L2* (**E**) and *L3* (**F**) fibrils with the β1 - β10 colored green or yellow and loops in gray. **G-I:** Overlay of a sharpened high-resolution map shown in blue (**G**), green (**H**), and orange (**I**) and an unsharpened, 4.5 Å low-pass filtered density is shown in gray. The backbone of the model is shown as black ribbon. Unsharpened densities highlighted with a yellow background are reminiscent of lipid micelles.

The *L1* fibrils were determined to a resolution of 3.2 Å for *L1A* (**Fig. S3A**) and 3.0 Å for *L1B* and *L1C* (**Fig. S3B, C**), allowing to accurately model residues M1-Q99 (**Fig. S4-S6**). Each monomer in the *L1* protofilament comprises ten β-strands (β1 to β10) with nine connecting loops. Strands β2 and β8 form the tightly packed core, stabilized by a predominant hydrophobic steric zipper (*15*).

While the *L1A* fibril consists of a single *L1* protofilament (**Fig. 1B**), *L1B* and *L1C* fibrils are composed of two identical and intertwined *L1* protofilaments (**Fig. 1C, D**). The *L1B* and *L1C* fibrils differ in their protofilament interfaces. In the *L1B* fibril, both protofilaments are related by an approximate 2_1_ screw symmetry and the protofilaments are tilted by ∼37° to each other (**Fig. S7**). Protofilament dimerization, mediated by hydrophobic interactions between residues M1 and V40 as well as G41, accommodates a wide cavity in the protofilament interface. The protofilaments in the *L1C* fibril, on the other hand, are related by C_2_ symmetry and ionic interactions between residues K45 and E46 form the inter-protofilament interface.

The *L2A* fibril was determined to a resolution of 2.7 Å (**Fig. 2B, S3D, S8**), 3.1 Å for *L2B* (**Fig. 2C, S3E, S9**), and 2.8 Å for *L3A* (**Fig. 2D, S3F, S10**). The *L2* fold is similar but not identical to αSyn “polymorph 2” (PDB-ID: 6SST), firstly reported by Guerrero-Ferreira *et al*. (*16*) (**Fig. 2E, S11A, B**). The main structural difference between the folds is a shift of strand β5 by about 10 Å with respect to strands β8 and β1. On the other hand, the *L3* fold is similar to the fold determined for the E46K variant (PDB-ID: 6UFR) (*17*) (**Fig. 2F, S11C**). Here, the main structural differences are shifts of strands β4-β7 relative to their position in 6UFR and the presence of the N-terminal _14_GVVAAA_19_, which forms another β-strand neighboring β7.

The *L2A* fibril is composed of three identical *L2* protofilaments related by a C_3_ symmetry. Interestingly, the three protofilaments are separated by ∼20 Å and thus show no direct protein-protein contacts (**Fig. 2B)**. In the *L2B* fibril, two identical but asymmetrically arranged *L2* protofilaments form the mature fibril and the *L3A* fibril reveals a similar protofilament orientation (**Fig. 2C**-**F**). In both *L2B* and *L3A* fibrils, the helical axes of the two protofilaments point in opposite directions, which leads to an identical structure and elongation kinetics of both fibril ends. These structures belong to a symmetry class of amyloid fibrils which has been postulated (*18*) but, so far, not been observed experimentally.

For all fibril structures, the cross-section cryo-EM maps reveal additional ring- and rod-shaped densities at the fibril surface (**Fig. 1F-H, 2G-I, 3A**), which together are reminiscent of the cross-section of phospholipid micelles. In addition, previous ssNMR experiments identified residues of αSyn that bind to phospholipids (*11*), and most of those are neighboring these non-proteinaceous densities (**Fig. 3A, B, D, F**). We, therefore, assign the micelle-like cross-section densities to phospholipids bound to the fibril surfaces and refer to these protein-lipid aggregates as lipidic fibrils.

**Fig. 3.**
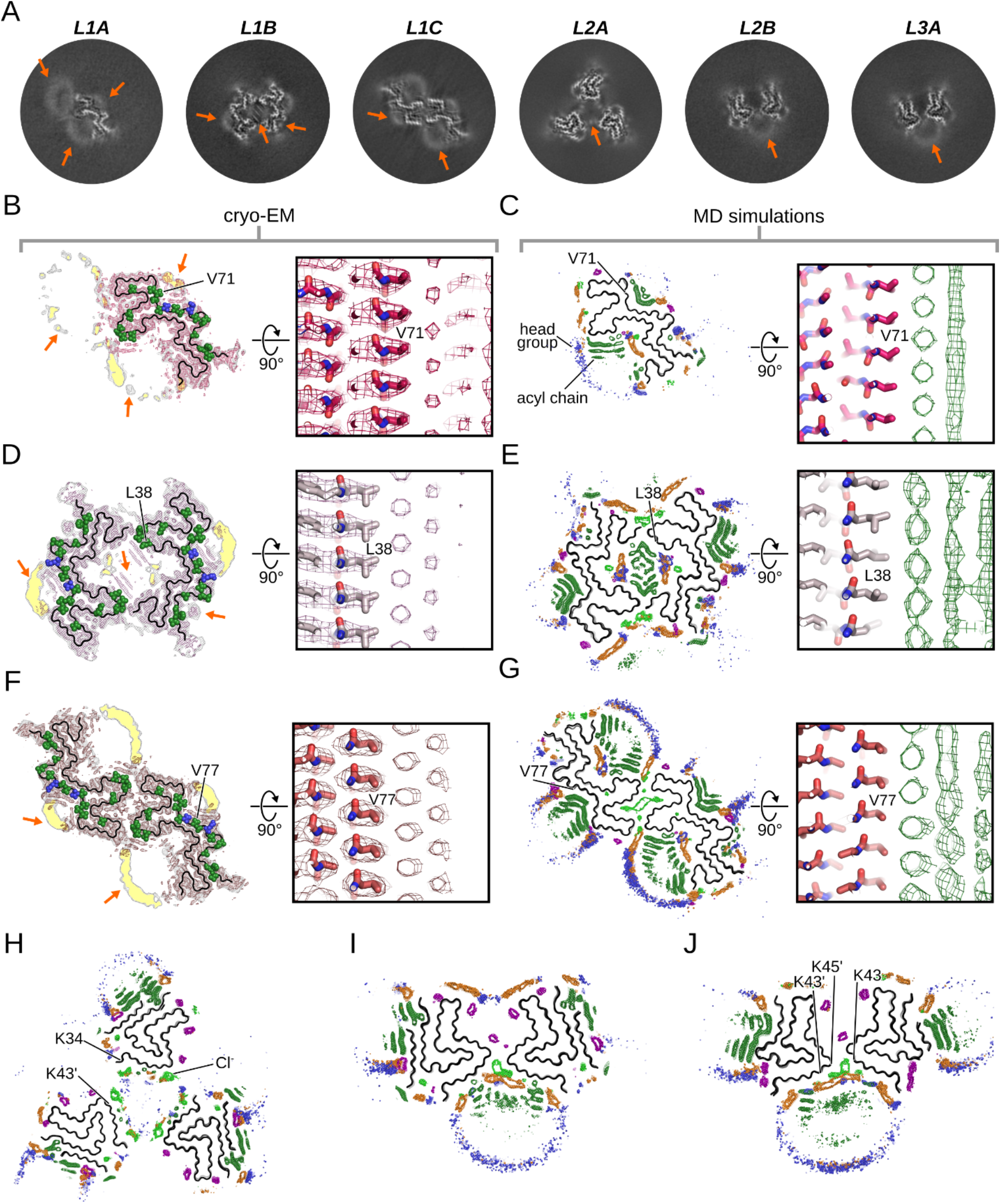
Fibril-lipid interactions. **A:** Central slice of the unsharpened refined cryo-EM maps. **B, D, F**: Superimposition of the reconstructed cryo-EM maps and the atomic model. Sharpened, high-resolution maps are shown in magenta (**B**, *L1A*), violet (**D**, *L1B*), and red (**F**, *L1C*). Unsharpened, 4.5 Å low-pass filtered density is shown in gray. The backbone of the model is shown as black ribbon, with residues binding to the acyl chain (green) or choline moiety (blue) of phospholipids shown as spheres (*11*). In **A+D**, the arrows highlight non-fibrillar densities. **C, E, G:** The grids indicate the probability density of the lipid acyl chain (dark green), phosphate (orange), and the choline nitrogen (blue), and sodium (purple), and chloride (light green) ions throughout MD simulations of the *L1A* (**C**), *L1B*, (**E**), and *L1C* (**G**) fibril. In **B-G**, the right panels show a close-up view visualizing the ordered packing of the lipid molecules along the helical axis. **H-J**: Probability density of the lipids throughout MD simulations of lipid diffusion for the *L2A* (**H**), *L2B*, (**I**), and *L3A* (**J**) fibril.

To validate this interpretation, we performed MD simulations of free lipid diffusion in the presence of the αSyn fibril structures determined here. Comparable simulations have successfully identified binding sites for biomarkers on other amyloid fibrils (*19, 20*). The initially randomly distributed lipid molecules associate towards micelle-like aggregates (**Movie S1**) and subsequently bind to predominantly hydrophobic areas on the fibril surface (**Fig. S12-S18**). The conversion of the SUVs used for the preparation of the lipidic fibrils to such small lipid aggregates upon fibril formation was confirmed by ^31^P ssNMR (**Fig. S19**). We calculated average density grids, showing the probability distribution of the lipids relative to the αSyn fibrils, averaged over multiple independent MD trajectories. Comparison to the cryo-EM cross-sections shows that the average lipid density from MD simulations almost perfectly matches the micelle-like densities in the cryo-EM cross-sections (**Fig. 3C, E, G**). Additionally, the averaged lipid density from MD simulations also reflects the periodical arrangement of the rod-shaped densities along the helical axis seen in the cryo-EM maps. Consequently, the MD data corroborates our assumption that the non-proteinaceous densities in the cryo-EM cross-sections likely are phospholipids bound to the fibril surface. In particular, the ring-shaped densities in the cryo-EM cross-sections likely constitute the lipid head groups, while the rod-shaped densities can be attributed to the lipid acyl chains. **Fig. 4** shows a POPC molecule modeled into the most well-defined of these densities.

**Fig. 4.**
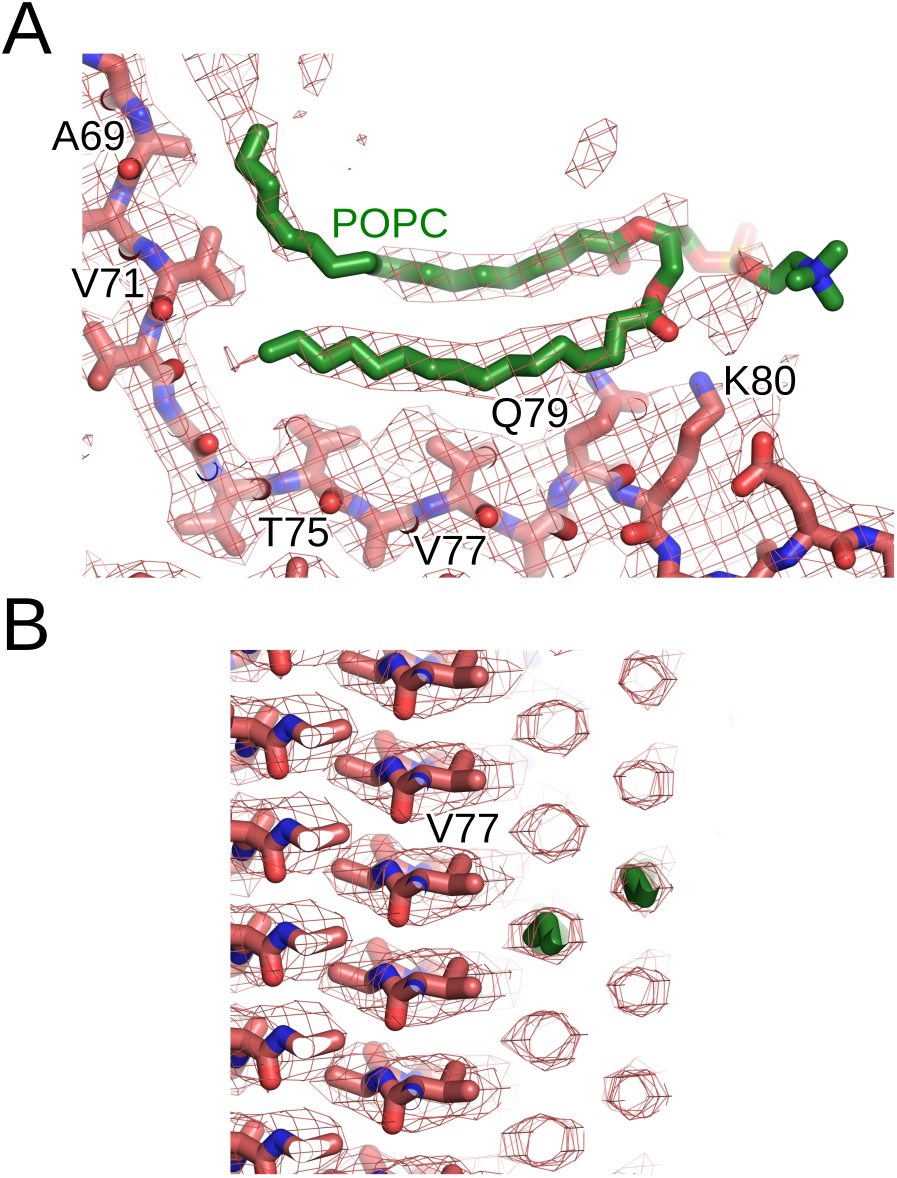
A fibril-lipid binding-mode model. One POPC molecule (green sticks) was modeled into the sharpened map of *L1C* and is shown in a view along the fibril axis (**A**) and perpendicular to the fibril axis (**B**).

The central cavity in the *L1B* fibril is occupied by lipids, with their head groups pointing towards charged residues K6, K21, and E20 and their acyl chains pointing towards the center. The cavity also harbors two areas likely occupied by chloride ions (Cl^-^), bridging interactions of positively charged residues K6 and K21 with the choline moiety of the lipids (**Fig. 3D, E**). For the *L1C* fibril, MD simulations revealed a high probability for Cl^-^ ions in the hydrophilic interface involving residues K43, K45, and E57 (**Fig. 3F, G**).

A striking feature of the *L2A* fibril is the bridging of lipid molecules that span the ∼20 Å gap between the protofilaments. Furthermore, head group densities of these lipids partially overlap with densities for Cl^-^ (**Fig. 3A, H**). Hence, the negatively charged phosphate groups, the positively charged choline moiety, and the Cl^-^ ions together form the bridge between K34 and K43 in the individual protofilaments by forming a well-ordered interaction network. This bridging is also seen for *L2B* and *L3A* fibrils, where Cl^-^ ions colocalize with head groups at the interface between K43 and K45 (**Fig. 3A, I, J**).

Although the *L2* and *L3* folds appear reminiscent of reported structures (*16, 17*), fibril-lipid interactions favor novel quaternary protofilament arrangements. In the *L2A* fibril, lipid-mediated interactions seem essential by connecting the neighboring protofilaments. Lipid-mediated interactions might also be responsible for the protofilaments pointing in opposite directions in the *L2B* and *L3A* fibrils, as in this configuration, the segment _34_KEGVLYVGSK_45_ in both protofilaments is in contact with the same phospholipid micelle.

While micelle-like lipid arrangements at the fibril surface can potentially also result from diffusion of lipid molecules after fibril formation, lipids in the central cavity of *L2A* seem less likely to originate from this process, as lipids mediate the interaction between the protofilaments, suggesting the presence of lipids already during fibril assembly. It is curious to note that the segment _35_EGVLYV_40_ in the lipidic *L1* fibrils or _34_KEGVLYVGSK_45_ in lipidic *L2* and *L3* fibrils are in contact with lipids, which suggests that this stretch of residues could play an important role throughout fibril formation. Indeed, residues within this sequence, such as Y39, have previously been identified to play a crucial role in lipid binding, aggregation kinetics, and function (*21, 22*).

Although the mechanisms of αSyn aggregation and toxicity *in vivo* are still under debate (*23, 24*), disruption of intracellular vesicles is one potential mechanism for the cellular toxicity mediated by αSyn fibrils (*25*) as well as oligomers (*26*). Reynolds *et al*. proposed a mechanism in which the abnormal aggregation of αSyn is linked to continuous lipid extraction mediated by the growing aggregates, eventually leading to membrane disruption (*27*). The finding of direct fibril-lipids interactions due to their lipid-associated aggregation may provide the structural basis for the proposed lipid extraction mechanism (*27*). In addition, the lipid-coated fibrils reported here give a structural rationale to the previously suggested lipid co-aggregation with αSyn fibrils (*28*).

In recent years, a growing number of *ex vivo* cryo-EM fibril structures have been discovered that are characteristic of different diseases (*29-31*). That none of these so far are lipidic fibrils might be explained by the use of detergent during the isolation of fibrils from patient tissue.

In conclusion, we report six fibril structures of αSyn, revealing how lipid molecules bind directly to the fibril surface. Insights obtained from these lipidic fibrils emphasize that studying αSyn aggregates in the presence of lipids is relevant for understanding the molecular basis of α-synucleinopathies. Furthermore, modulation of fibril-lipid interactions may also provide a promising strategy in searching for novel therapeutic interventions.

## Acknowledgments

BF, JAG, and GFS are grateful for computational support and infrastructure provided by the ‘‘Zentrum für Informations-und Medientechnologie’’ (ZIM) at the Heinrich Heine University Düsseldorf and the computing time provided by Forschungszentrum Jülich on the supercomputer JURECA/JURECA-DC at Jülich Supercomputing Centre (JSC). We thank Karin Giller and Melanie Wegstroth for excellent technical help with protein sample preparation.

## Author contributions

Conceptualization: CG, GFS

Protein expression and purification: SB

Preparation of fibril samples: LeA

ssNMR data acquisition and interpretation: LeA, EEN, LBA

Cryo-EM grid preparation and data collection: CD

Image processing, reconstruction and model building: BF

Visualization: BF, LeA

Project administration: BF, LeA, CG, GFS

Supervision: CG, GFS

Writing – original draft: BF, LeA, CG, GFS

Writing – review & editing: All authors

## Data and materials availability

Cryo-EM maps and motion-corrected micrographs have been deposited in the Electron Microscopy Data bank (EMDB) under the accession numbers EMD-XXX (*L1A*), EMD-XXX (*L1B*), EMD-XXX (*L1C*), EMD-XXX (*L2A*), EMD-XXX (*L2B*), and EMD-XXX (*L3A*). The corresponding atomic models have been deposited in the Protein Data Bank (PDB) under the accession numbers: XXXX (*L1A*), XXXX (*L1B*), XXXX (*L1C*), XXXX (*L2A*), XXXX (*L2B*), and XXXX (*L3A*). Additional data will be available from the corresponding authors upon reasonable request.

## Supplementary Materials

Materials and Methods

Figs. S1 to S14

Tables S1 to S2

Movies S1

## References

1. M. Goedert, Alzheimer’s and Parkinson’s diseases: The prion concept in relation to assembled amyloid-beta, tau, and alpha-synuclein. Science 349, 1255555 (2015).

2. M. Goedert, M. Masuda-Suzukake, B. Falcon, Like prions: The propagation of aggregated tau and alpha-synuclein in neurodegeneration. Brain 140, 266–278 (2017).

3. T. Uchihara, B. I. Giasson, Propagation of alpha-synuclein pathology: Hypotheses, discoveries, and yet unresolved questions from experimental and human brain studies. Acta Neuropathol. 131, 49–73 (2016).

4. M. G. Spillantini, R. A. Crowther, R. Jakes, M. Hasegawa, M. Goedert, alpha-synuclein in filamentous inclusions of Lewy bodies from Parkinson’s disease and dementia with Lewy bodies. Proc. Natl. Acad. Sci. U. S. A. 95, 6469–6473 (1998).

5. M. Baba et al., Aggregation of alpha-synuclein in Lewy bodies of sporadic Parkinson’s disease and dementia with lewy bodies. Am. J. Pathol. 152, 879–884 (1998).

6. K. Araki et al., Parkinson’s disease is a type of amyloidosis featuring accumulation of amyloid fibrils of alpha-synuclein. Proc. Natl. Acad. Sci. U. S. A. 116, 17963–17969 (2019).

7. S. H. Shahmoradian et al., Lewy pathology in Parkinson’s disease consists of crowded organelles and lipid membranes. Nat. Neurosci. 22, 1099–1109 (2019).

8. W. P. Gai et al., In situ and in vitro study of colocalization and segregation of alpha-synuclein, ubiquitin, and lipids in Lewy bodies. Exp. Neurol. 166, 324–333 (2000).

9. W. A. den Jager, Sphingomyelin in Lewy inclusion bodies in Parkinson’s disease. Arch. Neurol. 21, 615–619 (1969).

10. R. Stok, A. Ashkenazi, Lipids as the key to understanding alpha-synuclein behaviour in Parkinson disease. Nat. Rev. Mol. Cell Biol. 21, 357–358 (2020).

11. L. Antonschmidt et al., Insights into the molecular mechanism of amyloid filament formation: Segmental folding of alpha-synuclein on lipid membranes. Sci. Adv. 7, eabg2174 (2021).

12. S. Takamori et al., Molecular anatomy of a trafficking organelle. Cell 127, 831–846 (2006).

13. R. J. Perrin, W. S. Woods, D. F. Clayton, J. M. George, Interaction of human alpha-synuclein and Parkinson’s disease variants with phospholipids - Structural analysis using site-directed mutagenesis. J. Biol. Chem. 275, 34393–34398 (2000).

14. S. Kubo et al., A combinatorial code for the interaction of alpha-synuclein with membranes. J. Biol. Chem. 280, 31664–31672 (2005).

15. M. R. Sawaya, M. P. Hughes, J. A. Rodriguez, R. Riek, D. S. Eisenberg, The expanding amyloid family: Structure, stability, function, and pathogenesis. Cell 184, 4857–4873 (2021).

16. R. Guerrero-Ferreira et al., Two new polymorphic structures of human full-length alpha-synuclein fibrils solved by cryo-electron microscopy. Elife 8, (2019).

17. D. R. Boyer et al., The alpha-synuclein hereditary mutation E46K unlocks a more stable, pathogenic fibril structure. Proc. Natl. Acad. Sci. U. S. A. 117, 3592–3602 (2020).

18. D. S. Eisenberg, M. R. Sawaya, Structural studies of amyloid proteins at the molecular level. Annu Rev Biochem 86, 69–95 (2017).

19. B. Frieg, L. Gremer, H. Heise, D. Willbold, H. Gohlke, Binding modes of thioflavin T and Congo red to the fibril structure of amyloid-beta(1-42). Chem. Commun. 56, 7589–7592 (2020).

20. C. König et al., Binding sites for luminescent amyloid biomarkers from non-biased molecular dynamics simulations. Chem. Commun. 54, 3030–3033 (2018).

21. L. Fonseca-Ornelas et al., Small molecule-mediated stabilization of vesicle-associated helical alpha-synuclein inhibits pathogenic misfolding and aggregation. Nat. Commun. 5, 5857 (2014).

22. C. P. A. Doherty et al., A short motif in the N-terminal region of alpha-synuclein is critical for both aggregation and function. Nat. Struct. Mol. Biol. 27, 249–259 (2020).

23. Y. C. Wong, D. Krainc, alpha-synuclein toxicity in neurodegeneration: mechanism and therapeutic strategies. Nat. Med. 23, 151–163 (2017).

24. P. Alam, L. Bousset, R. Melki, D. E. Otzen, alpha-synuclein oligomers and fibrils: a spectrum of species, a spectrum of toxicities. J. Neurochem. 150, 522–534 (2019).

25. W. P. Flavin et al., Endocytic vesicle rupture is a conserved mechanism of cellular invasion by amyloid proteins. Acta Neuropathol. 134, 629–653 (2017).

26. G. Fusco et al., Structural basis of membrane disruption and cellular toxicity by alpha-synuclein oligomers. Science 358, 1440–1443 (2017).

27. N. P. Reynolds et al., Mechanism of membrane interaction and disruption by alpha-synuclein. J. Am. Chem. Soc. 133, 19366–19375 (2011).

28. E. Hellstrand, A. Nowacka, D. Topgaard, S. Linse, E. Sparr, Membrane lipid co-aggregation with alpha-synuclein fibrils. PLoS One 8, e77235 (2013).

29. A. W. P. Fitzpatrick et al., Cryo-EM structures of tau filaments from Alzheimer’s disease. Nature 547, 185–190 (2017).

30. M. Schweighauser et al., Structures of alpha-synuclein filaments from multiple system atrophy. Nature, (2020).

31. Y. Yang et al., Cryo-EM structures of amyloid-beta 42 filaments from human brains. Science 375, 167–172 (2022).

32. A. Mori, Y. Imai, N. Hattori, Lipids: key players that modulate alpha-synuclein toxicity and neurodegeneration in Parkinson’s disease. Int. J. Mol. Sci. 21, 3301 (2020).

