## Supplemental Material for "The 3D structure of lipidic fibrils of α-synuclein"

**This PDF file includes:**

Materials and Methods
Figs. S1 to S14
Tables S1 to S2
Captions for Movies S1

**Other Supplementary Materials for this manuscript include the following:**

Movies S1

### Materials and Methods

Protein expression and purification

αSyn was expressed recombinantly in *E. coli* strain BL21(DE3) and purified as described previously (*29*). Briefly, the protein was expressed in minimal medium at 37 °C. Cells were harvested 6h after induction, lysed by freeze-thaw cycles followed by sonication, boiled for 15 minutes and centrifuged at 48.000 x g for 45 minutes. From the supernatant DNA was precipitated with streptomycin (10 mg/ml) while stirring the ice-cold solution. After centrifugation αSyn was precipitated from the supernatant by adding ammonium sulfate to 0.36 g/ml. After another centrifugation step the pellet was resuspended in 25 mM Tris/HCl, pH 7.7 and the protein was further purified by anion exchange chromatography on a 30 ml POROS HQ column (PerSeptive Biosystems). To prepare monomeric αSyn without any aggregates, the protein was dialyzed against PBS buffer, pH 7.4, centrifuged at 106,000 x g for 1h at 4°C and filtrated through 0.22 µm ULTRAFREE-MC centrifugal filter units (Merck Millipore). The final protein concentration was adjusted to 0.33 mM.

Preparation of αSyn fibrils

Samples of αSyn fibrils were prepared as previously reported (*30*) In brief, vesicles were prepared by mixing 1-palmitoyl-2-oleoyl-*sn*-glycero-3-phosphocholine (POPC), 1-palmitoyl-2-oleoyl-*sn*-glycero-3-phosphate (POPA, sodium salt) dissolved in chloroform respectively and evaporating the solvent under a $N_2$-stream followed by lyophilization overnight. SUVs were obtained by repeated sonication of a solution of 1.5 mM POPC, 1.5 mM POPA. Vesicles were incubated with 70 µM $^{13}$C, $^{15}$N-labelled αS in buffer (50 mM HEPES, 100 mM NaCl, pH 7.4) at a lipid to protein ratio of 5:1 and subjected to repeated cycles of 30 s sonication (20 kHz) at 37 °C followed by an incubation period of 30 min. After 24 h (dataset 1), 48 h (dataset 2) and 20 h (dataset 3) respectively the samples were transferred to a Multitron incubator (Infors HT, Bottmingen, CH) and shaken at 100 rpm (50 mm throw) at 37 °C until a combined aggregation time of 96 h was reached. Aggregation was monitored regularly by mixing 5 µL of the aggregate solution with 2 mL of Thioflavin T containing buffer (100 µM ThT, 50 mM Glycine, pH 8.5) and measuring the fluorescence emission intensity at 482 nm in a Varian Cary Eclipse fluorescence spectrometer.

For Cryo-EM samples 700 µL of aggregate solution were then centrifuged for 5 min at 14.000 rpm in a F-45-18-11 Rotor in a 5418 R tabletop centrifuge (Eppendorf, Hamburg, GER). If fibrils did not pellet right away, the procedure was repeated until a visible pellet was obtained. The supernatant was removed and 50 µL of fresh buffer (5 mM HEPES, pH 7.4) were added and thoroughly mixed with the pellet to obtain a highly concentrated fibril solution.

For ssNMR samples a minimum of 1500 µL of the aggregate solution were centrifuged at 55.000 rpm (TLA-100.3 rotor in an Optima™ MAX-TL) for 1 h at 4 °C. After removal of the supernatant, samples were washed with fresh buffer (5 mM HEPES, pH 7.4) and subsequently centrifuged (10 min, 65.000 rpm, 18 °C). Excess moisture was carefully removed, and samples were packed into either 1.3 mm or 3.2 mm ssNMR rotors by cutting off the bottom of the tube and centrifuging the pellet directly into the rotor of choice through a custom-made filling device made from a truncated pipette tip. Finally, the sample was centrifuged into the rotor in an ultracentrifuge packing device for 30 min at 24.000 rpm in a SW 32 Ti rotor in an Optima™ L-80 XP Ultracentrifuge (both Beckman Coulter) (*31*).

ssNMR

3D (H)CANH experiments (*32*) $^{13}C$, $^{15}N$-labelled αS on an 800 MHz Bruker Avance III HD spectrometer at a magnetic field of 18.8 T or a 1200 MHz Bruker Avance NEO spectrometer at a magnetic field of 28.2 T each equipped with a 1.3 mm magic-angle spinning (MAS) HCN probe and MAS at 55 kHz. The temperature of the cooling gas was set to 250 K, resulting in an estimated sample temperature of 20 °C.

2D (H)NCA spectra were acquired on an 850 MHz Avance III spectrometer with a 3.2 mm MAS HCN probe at a magnetic field of 20.0 T and MAS at 17 kHz. The temperature of the cooling gas was set to 265 K, resulting in an estimated sample temperature of 20 °C.

$^{1}H$ decoupled $^{31}P$ spectra were acquired on an 600 MHz Avance III spectrometer with a 1.3 mm MAS HCN probe (equipped with a range coil for $^{31}P$ tuning) at a magnetic field of 14.1 T without MAS. The temperature of the cooling gas was set to 278.2 K and 310.2 K, resulting in estimated sample temperatures of 7 °C and 37 °C respectively. For spectra of vesicles, SUVs were prepared as described above. The resulting solution was lyophilized and resuspended in drops buffer (10 mM HEPES) to increase concentration. The resulting gel was centrifuged into the rotor in an ultracentrifuge packing device as described above.

Cryo-EM grid preparation and imaging

For cryo-EM grid preparation, 1.5 µL of fibril solution were applied to freshly glow-discharged R2/1 holey carbon film grids (Quantifoil). After the grids were blotted for 12 seconds at a blot force of 10, the grids were flash frozen in liquid ethane using a Mark IV Vitrobot (Thermo Fisher).

Cryo-EM data sets were collected on a Titan Krios transmission-electron microscope (Thermo Fisher) operated at 300 keV accelerating voltage and a nominal magnification of 81,000 x using a K3 direct electron detector (Gatan) in non-superresolution counting mode, corresponding to a calibrated pixel size of 1.05 Å. A total of 11,740, 7,836 and 7,744 movies were collected with SerialEM (*33*) for Datasets 01, 02 and 03, respectively. Movies of Dataset 01 were recorded over 50 frames accumulating a total dose of ~51 $e^{-}/A^{2}$, whereas movies of Dataset 02 and 03 contained 40 frames with a total dose of ~43 $e^{-}/A^{2}$. The range of defocus values collected spans from -0.5 µm to -2.0 µm. Collected movies were motion corrected and dose weighted on-the-fly using Warp (*34*).

Helical reconstruction pf αSyn fibrils

αSyn fibrils were reconstructed using RELION-3.1 (*35*), following the helical reconstruction scheme (*36*). Firstly, estimation of contrast transfer function parameters for each motion-corrected micrograph was performed using CTFFIND4 (*37*). For filament picking, we only considered micrographs with an estimated resolution of ≤ 3.8 Å (Dataset 01), ≤ 4.0 Å (Dataset 02), and ≤ 5.0 Å (Dataset 03) respectively (Tab. S1).

For 2D classification, we extracted particle segments using a box size of 600 pix downscaled to 200 pix and an inter-box distance of 13 pix (1.05 Å/pix). *L1A*, *L1B*, *L1C*, *L2A* fibrils were successfully separated at this 2D classification stage, whereas *L2B* and *L3A* were too similar on the 2D level.

For 3D classification, the classified segments after 2D classification were (re-)extracted using a box size of 250 pix and without downscaling. Starting from featureless cylinder filtered to 60 Å, several rounds of refinements were performed while progressively increasing the reference model's resolution. The helical rise was initially set to 4.75 Å and the twist was estimated from the micrographs. Once the β-strands were separated along the helical axis, we optimized the helical parameters (final parameters are reported in in Tab. S1). During 3D classification, we successfully separated *L2B* and *L3A* fibrils, which were then treated individually. We performed multiple rounds of 3D auto-refinement from here on until no further improvement of the map was observed. Standard RELION post-processing with a soft-edged solvent mask that includes the central 10 % of the box height yielded post-processed maps (*B*-factors are reported in Tab. S1). The resolution was estimated from the value of the FSC curve for two independently refined half-maps at 0.143 (Fig. S3). The optimized helical geometry was then applied to the post-processed maps yielding the final maps used for model building. For all fibrils, a left-handed twist was assumed.

Atomic model building and refinement

The atomic models of *L1* fibrils were built *de novo* in Coot (*38*). For *L2* fibrils, one protein chain was extracted from PDB-ID 6SST (*39*) of wild-type αSyn and used as the initial model. For *L3* fibrils, one protein chain from PDB-ID 6UFR (*40*) of E46K αSyn was extracted and used as the initial model. To the latter, the amino acid sequence was converted to wild-type αSyn (UniProt: P37840) and the N-terminal region G14 to A19 was built *de novo* in Coot (*38*). Subsequent refinement in real space was conducted using PHENIX (*41, 42*) and Coot (*38*) in an iterative manner. The resulting models were validated with MolProbity (*43*) and details about the atomic models are described in Tab. S2.

To visualize the lipid interactions, we used the sharpened *L1C* map and initially modeled a POPC molecule into the density, again using Coot (*38*). Subsequently, another round of real space refinement was conducted using PHENIX (*41, 42*).

Molecular dynamics simulations of lipid diffusion

To investigate where and how the lipids interact with the different types of αSyn fibrils, we performed unbiased molecular dynamics (MD) simulations of POPC and POPA in the presence of the αSyn fibrils. A filament was always composed of 20 helically arranged peptide chains. Except for residue M1 in *L1* fibrils, ACE- and NME-caps were connected to the N- and C-termini, respectively, to avoid artificially charged termini.

We then used PACKMOL (*44*) to, first, center the αSyn fibril in a rectangular simulation box, and, second, to randomly place POPC and POPA lipids, sodium ($Na^+$) and chloride ($Cl^-$) ions, and water molecules around the αSyn fibril. We added additional $Na^+$ or $Cl^-$ counter ions to enforce the neutrality of the systems. In the final setup, we mimicked the experimental conditions used for αSyn fibril aggregation (*30*), meaning that side chains are prepared for pH 7.4, the NaCl concentration is 100 mM, and a molar lipid/protein ratio is 10 (ratio of 1:1 for the lipids).

The Amber ff19SB force field (*45*) was applied to describe the αSyn fibrils and the Lipid17 force field (*46*) to describe the POPC and POPA molecules. Ion Parameters for monovalent ions were taken from ref. (*47*) and used in with the OPC water model (*48*).

The exact minimization, thermalization (towards 300 K), and density adaptation (towards 1 g/cm$^3$) protocol is reported in ref. (*49*), which was applied previously to study ligand binding processes to amyloid fibrils (*50*). The conformations after thermalization and density adaptation served as starting points for subsequent NPT production simulations. Therefore, the initial velocities were randomly assigned during the first step of the following NPT production simulation, such that each simulation can be considered as an independent replica. For each αSyn fibril, we completed eight independent NPT production simulations at 300 K and 1 bar for 1 µs each. Importantly, we restrained the backbone to the initial atomic coordinates. However, all other molecules, including POPC and POPA, were allowed to diffuse freely and we did not apply any artificial guiding force. During production simulations, Newton's equations of motion were integrated in 4 fs intervals, applying the hydrogen mass repartitioning approach (*51*) to all non-water molecules, which were handled by the SHAKE algorithm (*52*). Coordinates were stored into a trajectory file every 200 ps. The minimization, thermalization, and density adaptation were performed using the pmemd.MPI (*53*) module from Amber20/AmberTools21 (*54*), while the production simulations were performed with the pmemd.CUDA module (*55*).

Determination of the binding region for lipids

We used *cpptraj* (*56*) from Amber20/AmberTools21 (*54*) to calculate 3D density grids (normalized to the number of considered conformations) separately for the lipids' acyl chain, the phosphate atom, and the choline nitrogen atom. These grids represent the probability density of a molecule position relative to the centered fibril structure. Initially, we calculated the 3D density grids for each trajectory, constantly increasing the time range for the analysis in 0.1 μs intervals. Thereby, we observed only minimal changes when extending the analysis time from 0.9 μs ns to 1.0 μs, such that we assumed converged distributions of the lipid molecules. Hence, the average density grids were calculated over all conformations of the 0.9 μs to 1.0 μs interval of all MD simulations replicates.

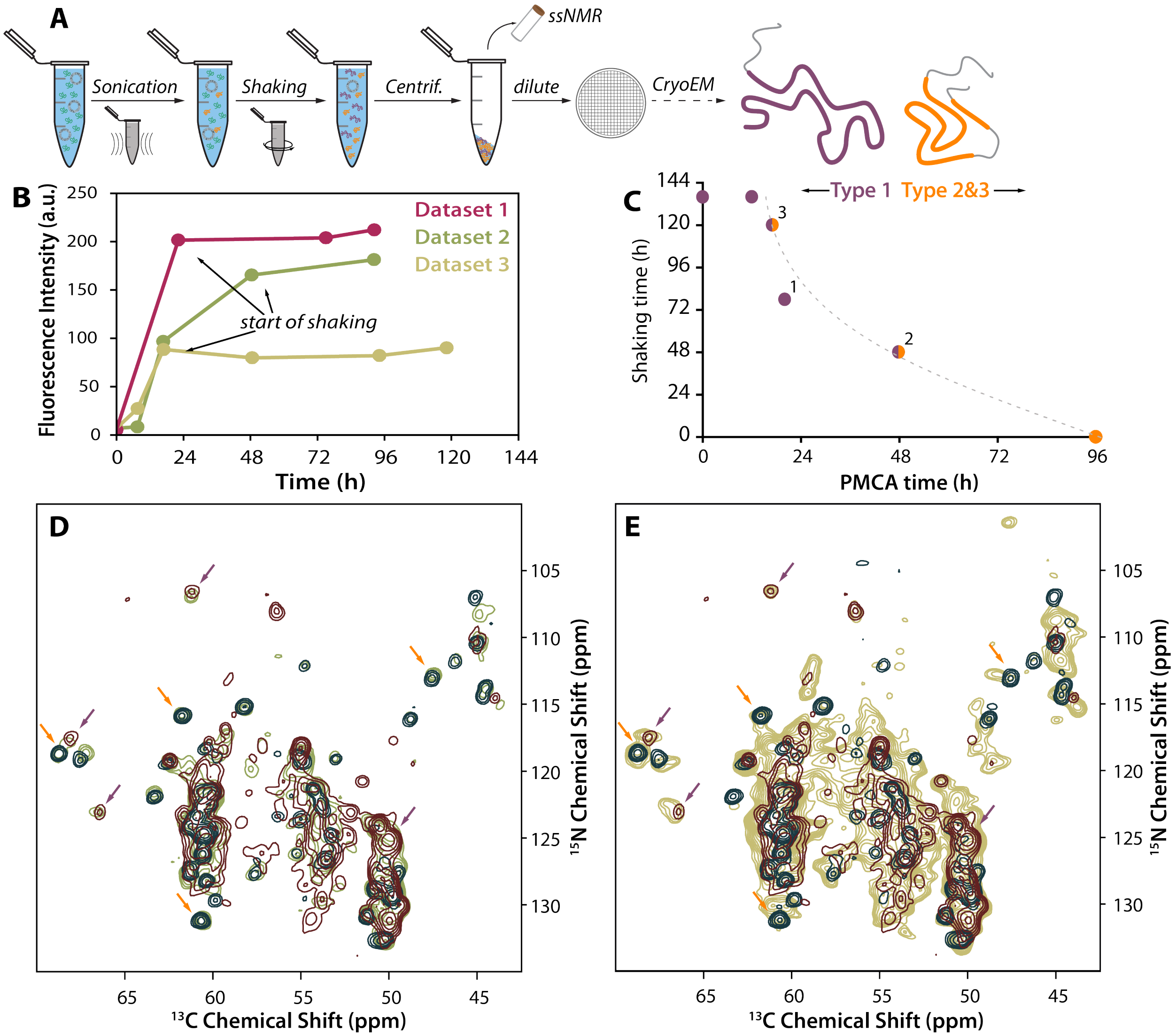


**Fig. S1. Preparation workflow and ssNMR analysis of fibril samples**

**A:** Workflow schematic for preparation of αSyn fibrils in this study. **B:** ThT fluorescence data of individual samples analyzed by cryo-EM. **C**: Correlation plot of times spent under different agitation conditions. Points are color coded by the dominant fibril types. Characterization of type 1 (purple) and type 2 (orange) was done by ssNMR (fibril subtypes were indistinguishable) and in labelled cases by Cryo-EM (datasets 1-3). **D**: (H)CANH spectra of αS fibrils used for dataset 1 acquired at 800 MHz with 55 kHz MAS (green) and **E**: (H)NCA of αS fibrils used for dataset 2 acquired at 850 MHz with 17 kHz MAS (yellow)compared to spectra of fibrils prepared under purely PMCA (blue, 950 MHz, 100 kHz MAS) and shaking conditions (red, 1200 MHz, 55 kHz MAS). Arrows indicate characteristic peaks originating from either *L1* (purple) or *L2*/*L3* fibrils (orange), showing that in either sample a mixture of both fibril types is present. Spectra of fibrils prepared under PMCA conditions (blue) are reproduced from ref. (*30*).

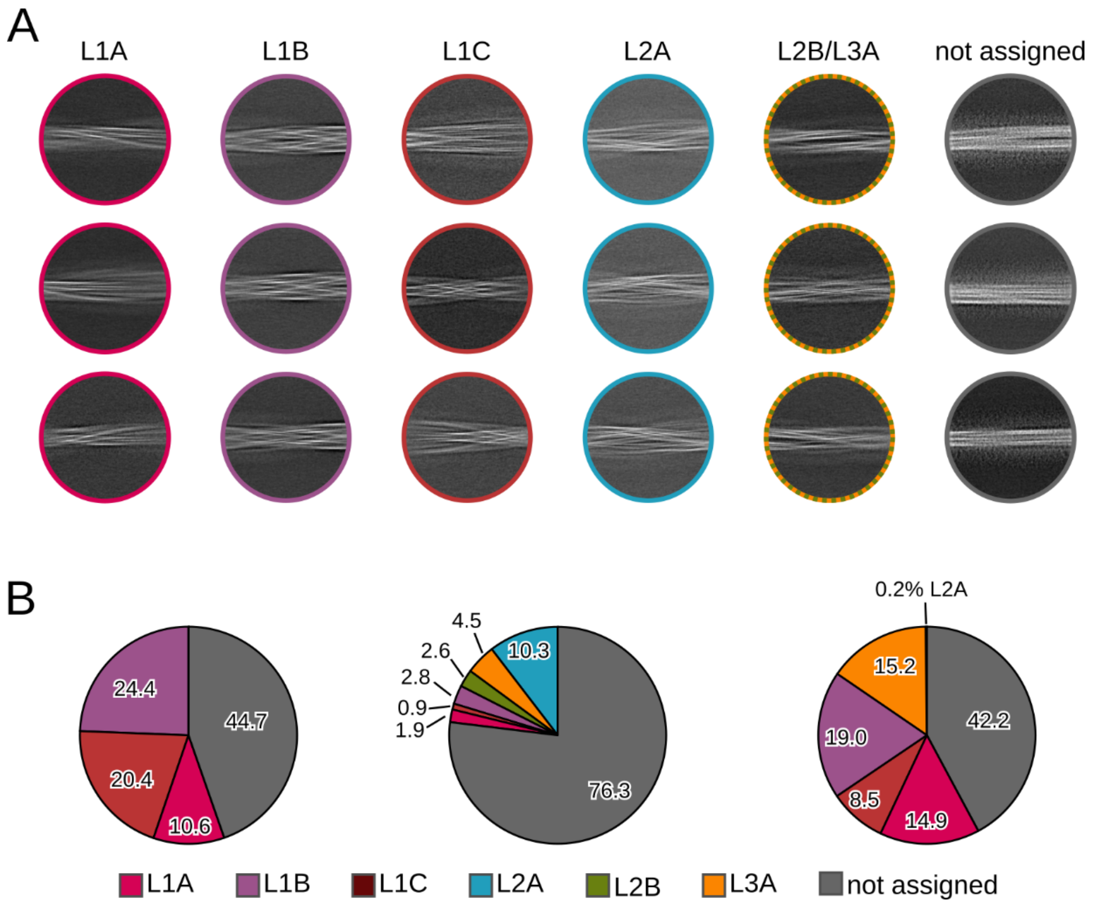


**Fig. S2. 2D class averages and polymorph distribution**

**A**: Representative 2D class averages for all lipid-induced αSyn fibrils and segments that could not be assigned to any of the polymorphs after 2D classification, due to the lack of well-defined and clear filament features. Instead, the unassigned classes are not sharp and partially very fuzzy at the fibril surface. **B**: Pie charts visualizing the relative population (labels in %) of each lipid-induced αSyn fibril polymorphs in dataset 01 (left) 02, (middle), and 03 (right).

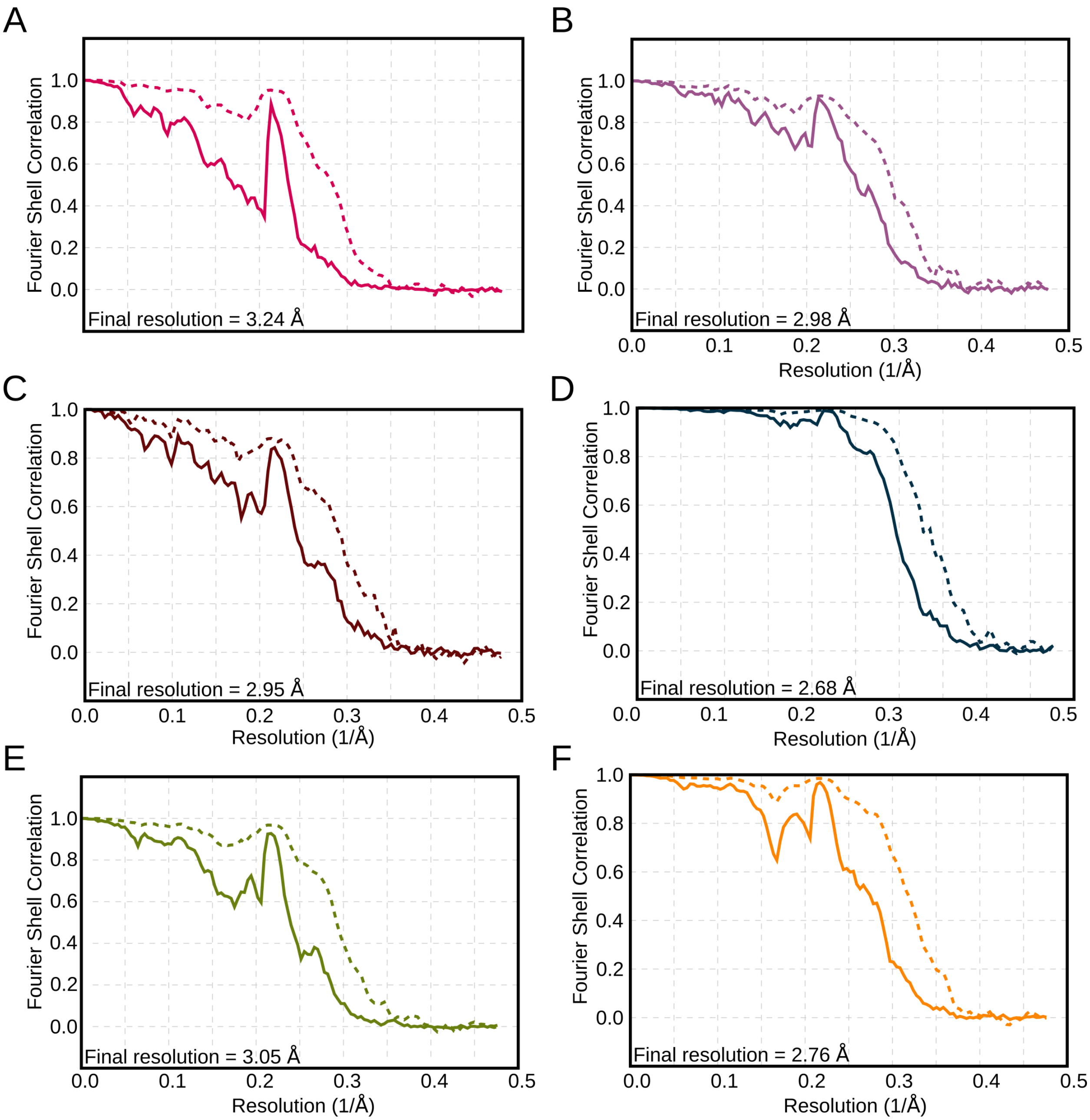


**Fig. S3. Fourier shell correlation curves.**
Fourier shell correlation (FSC) curves for *L1A* (**A**), *L1B* (**B**), *L1C* (**C**), *L2A* (**D**), *L2B* (**E**), and *L3A* (**F**). FSC curves are shown for two independently refined unmasked (solid lines) and masked (dashed lines) half-maps. The z-percentage is 0.1 all cases. The final resolution is shown in the plot and was estimated from the value of the FSC curve for two independently refined masked half-maps at 0.143.

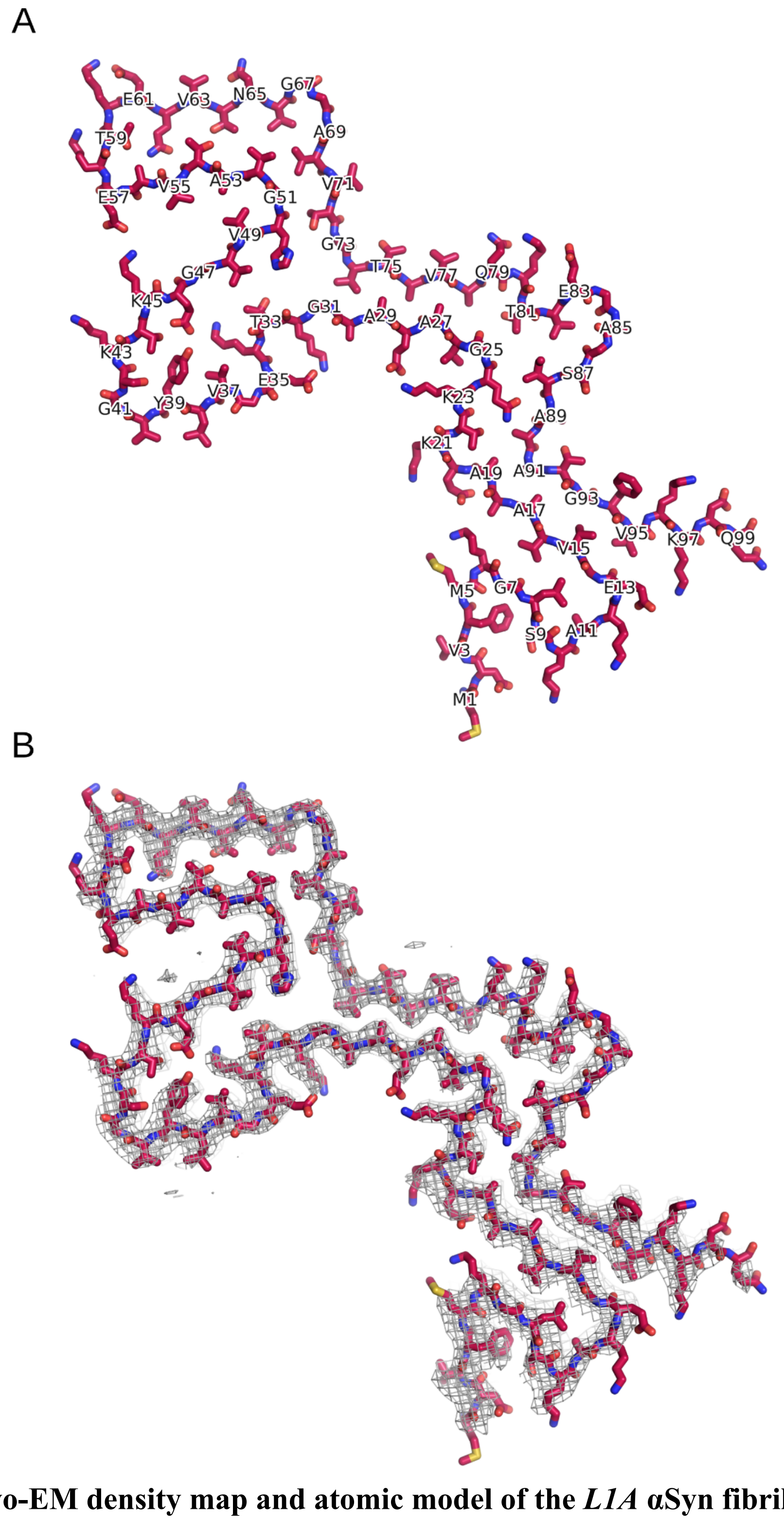


**Fig. S4. The cryo-EM density map and atomic model of the *L1A* αSyn fibril.**

**A**: The atomic model of the *L1A* αSyn fibril shown as stick model. For clarity, only every second amino acid is labeled. **B**: Superposition of the atomic model (shown in **A**) and the central slice of the density map with a width of 10.5 Å (10 pixel, 1.05 Å/pixel; gray isomesh; contour level of 0.05). Due to the tilt in the z-direction the atomic model is only partially visible in the central slice.

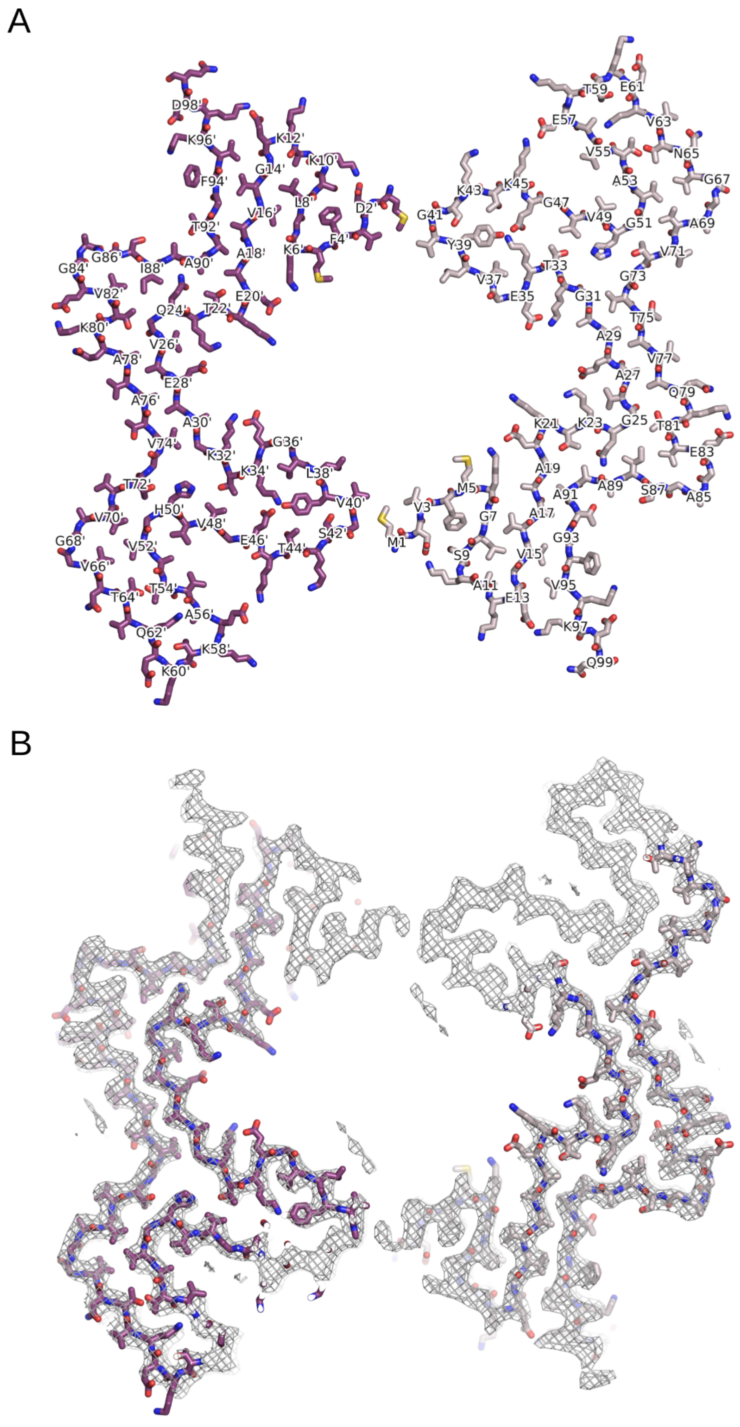


**Fig. S5. The cryo-EM map and atomic model of the *L1B* αSyn fibril.**

**A**: The atomic model of the L1B αSyn fibril shown as stick model. The two protofilaments are colored in different shades of purple. Even and odd numberings are given on one protofilament each. Amino acids from the darker colored protofilament are labeled with an additional prime. **B**: Superposition of the atomic model (shown in **A**) and the central slice of the density map with a width of 10.5 Å (10 pixel, 1.05 Å/pixel; gray isomesh; contour level of 0.06). Due to the tilt in the z-direction the atomic model is only partially visible in the central slice.

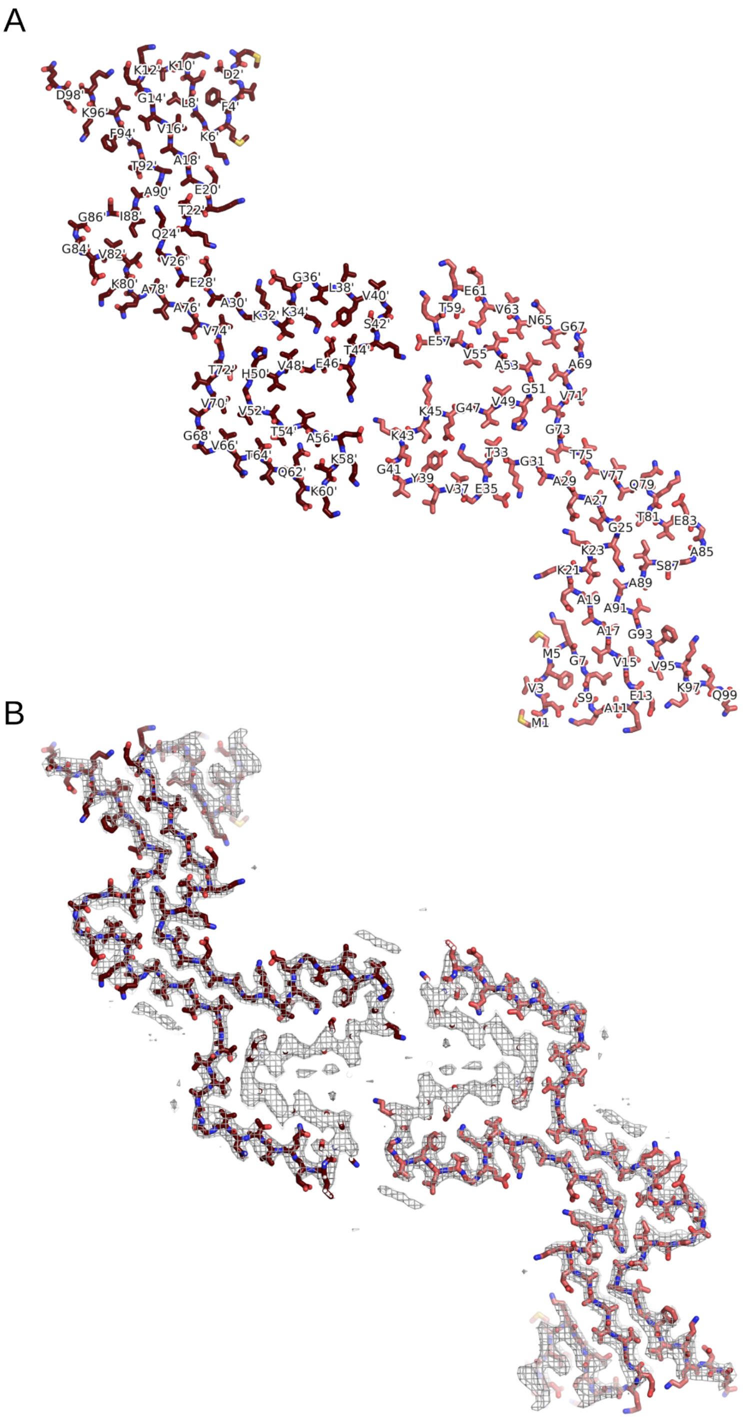


**Fig. S6. The cryo-EM density map and atomic model of the *L1C* αSyn fibril.**

**A**: The atomic model of the *L1C* αSyn fibril shown as stick model. The two protofilaments are colored in different shades of red. Even and odd numberings are given on one protofilament each. Amino acids from the darker colored protofilament are labeled with an additional prime. **B**: Superposition of the atomic model (shown in **A**) and the central slice of the density map with a width of 10.5 Å (10 pixel, 1.05 Å/pixel; gray isomesh; contour level of 0.05). Due to the tilt in the z-direction the atomic model is only partially visible in the central slice.

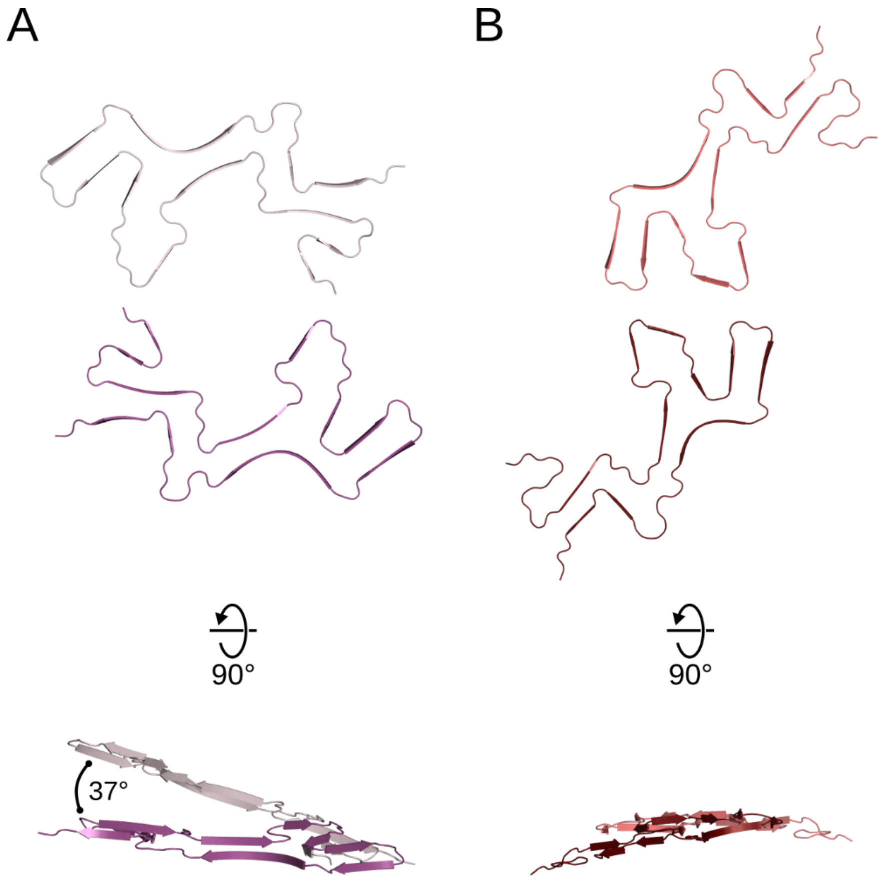


**Fig. S7. Pronounced intertwining of protofilaments in the *L1B* fibril.**

Two central protein chains extracted from the *L1B* (**A**) and *L1C* (**B**) αSyn fibril models in top and side-view.

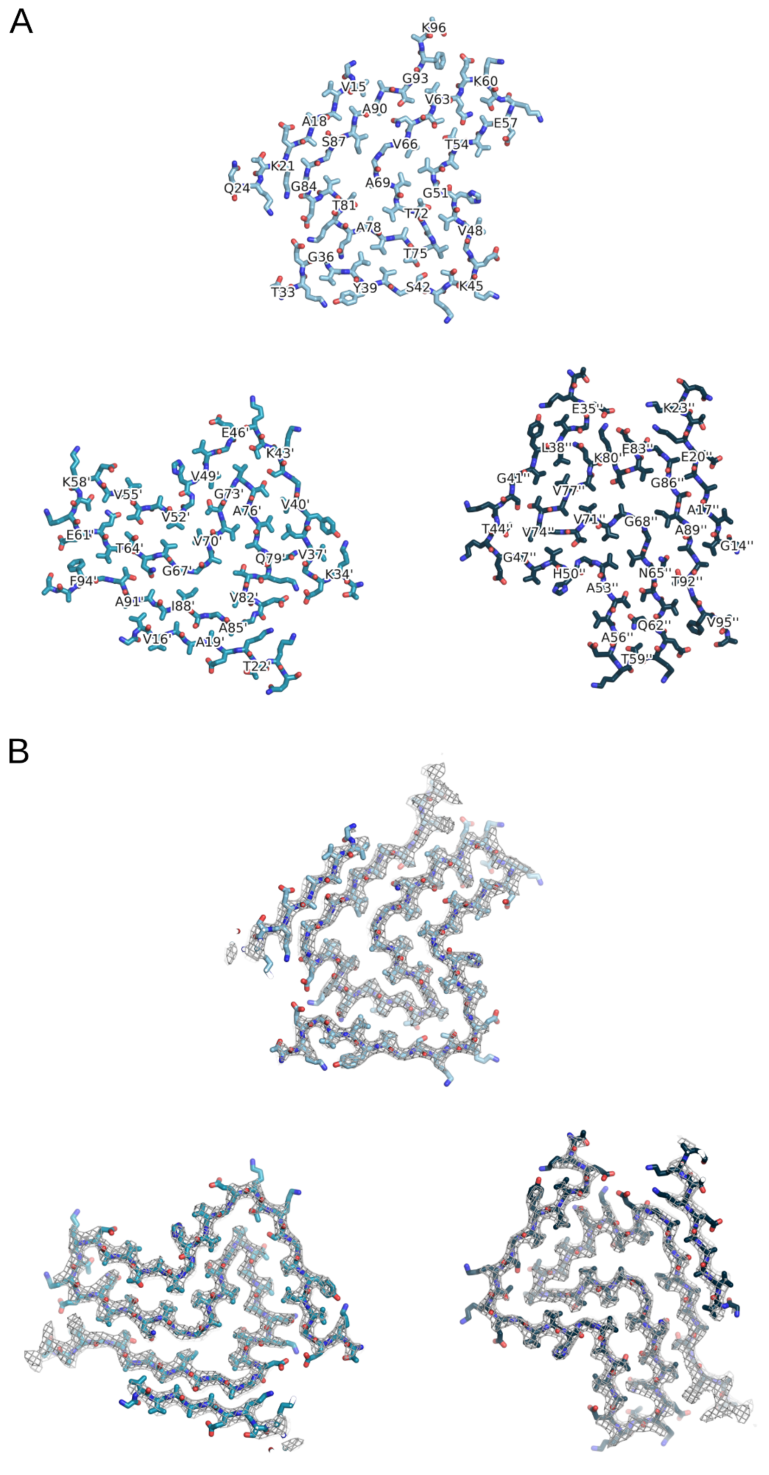


**Fig. S8. The cryo-EM map and atomic model of the *L2A* αSyn fibril.**

**A**: The atomic model of the *L2A* αSyn fibril shown as a stick model. The two protofilaments are colored in different shades of blue. Numberings are given on one protofilament each. Amino acids from the darker colored protofilaments are labeled with one or two additional primes. **B**: Superposition of the atomic model (shown in **A**) and the central slice of the density map with a width of 10.5 Å (10 pixel, 1.05 Å/pixel; gray isomesh; contour level of 0.076). Due to the tilt in the z-direction the atomic model is only partially visible in the central slice.

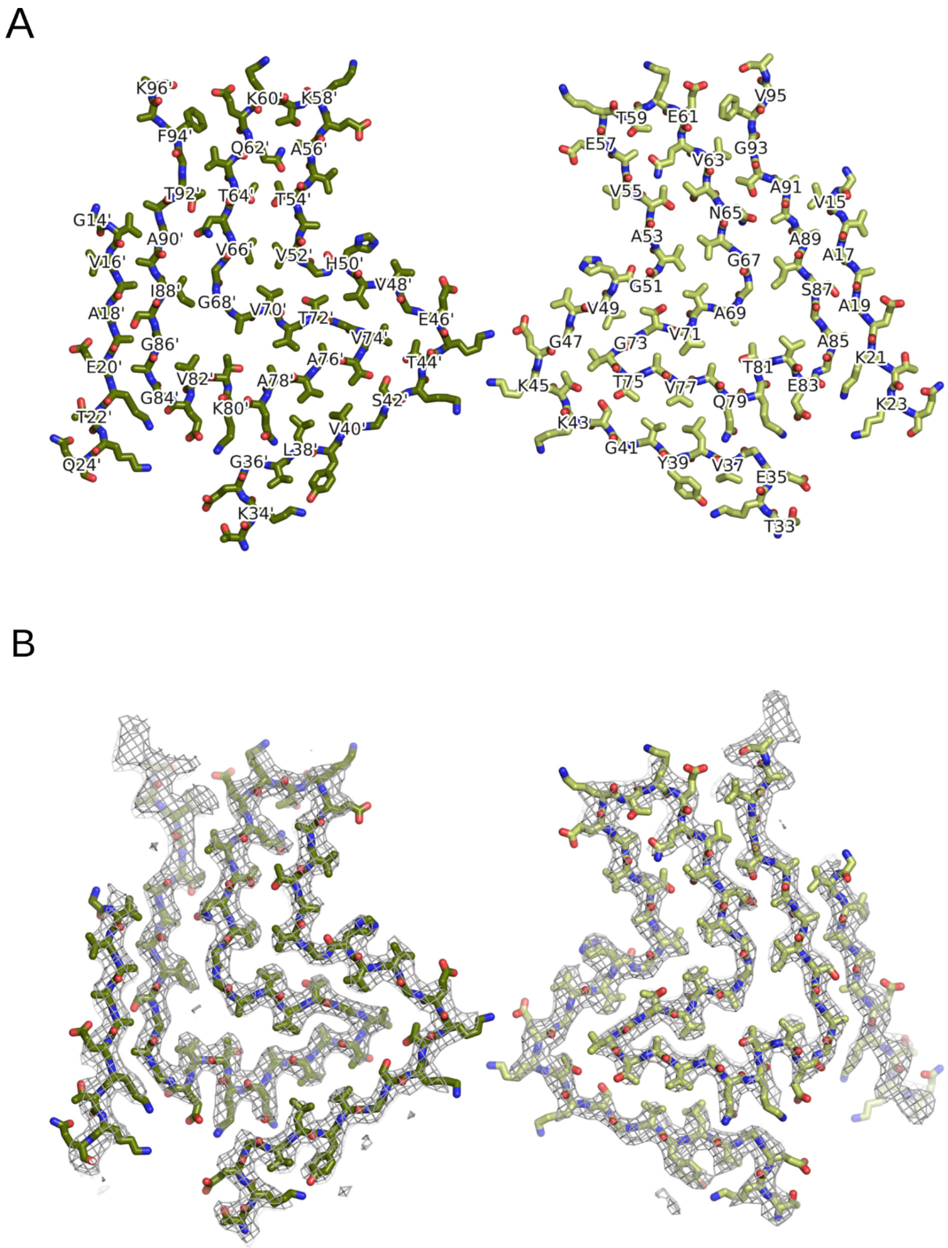


**Fig. S9. The cryo-EM map and atomic model of the *L2B* αSyn fibril.**
**A**: The atomic model of the *L2B* αSyn fibril shows as stick model. The two protofilaments are colored in different shades of green. Even and odd numberings are given on one protofilament each. Amino acids from the darker colored protofilament are labeled with an additional prime. **B**: Superposition of the atomic model (shown in **A**) and the central slice of the density map with a width of 10.5 Å (10 pixel, 1.05 Å/pixel; gray isomesh; contour level of 0.0519). Due to the tilt in the z-direction the atomic model is only partially visible in the central slice.

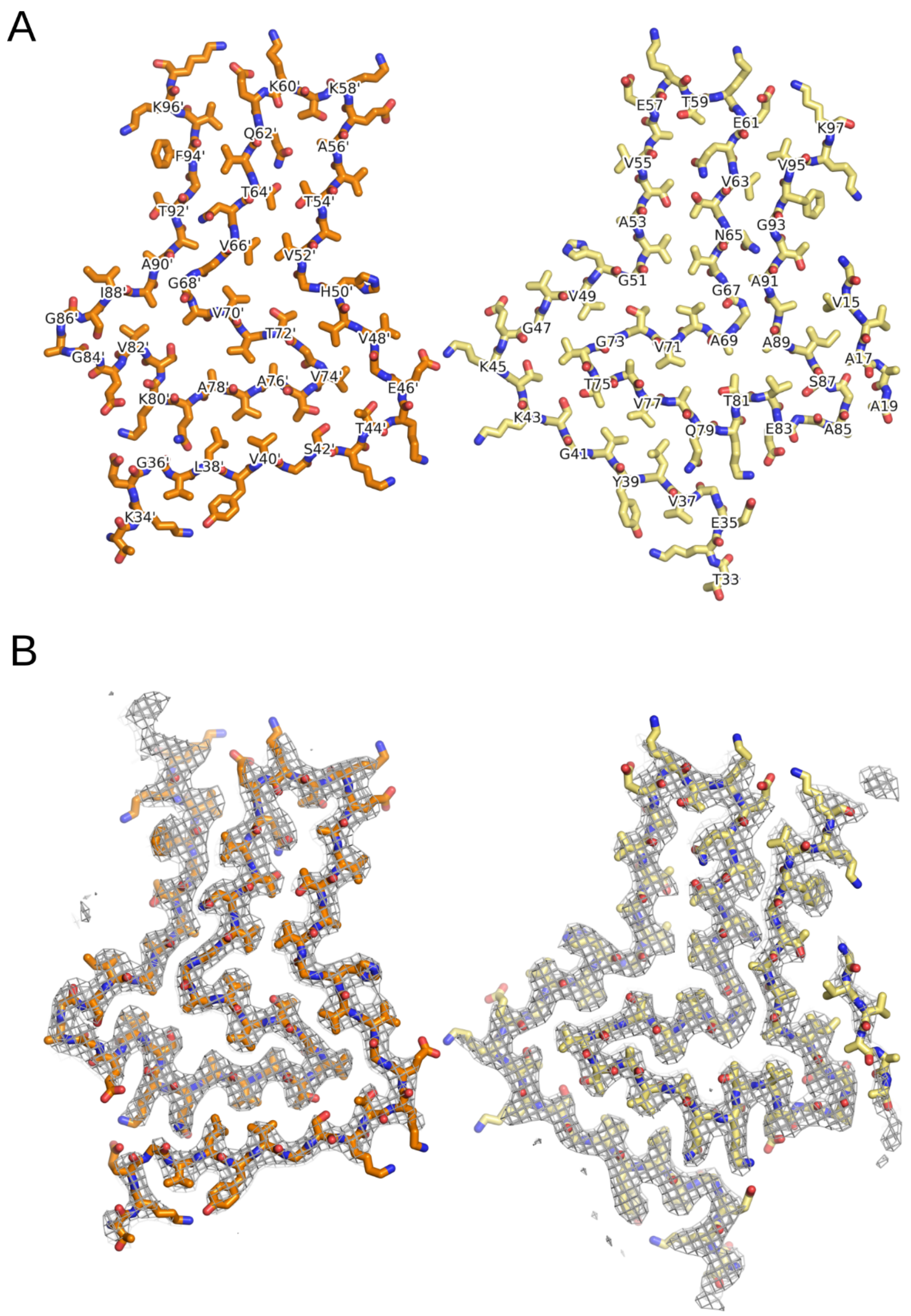


**Fig. S10. The cryo-EM map and atomic model of the *L2B* αSyn fibril.**

**A**: The atomic model of the *L2B* αSyn fibril shown as a stick model. The two protofilaments are colored in different shades of orange. Even and odd numberings are given on one protofilament each. Amino acids from the darker colored protofilament are labeled with an additional prime. **B**: Superposition of the atomic model (shown in **A**) and the central slice of the density map with a width of 10.5 Å (10 pixel, 1.05 Å/pixel; gray isomesh; contour level of 0.0512). Due to the tilt in the z-direction the atomic model is only partially visible in the central slice.

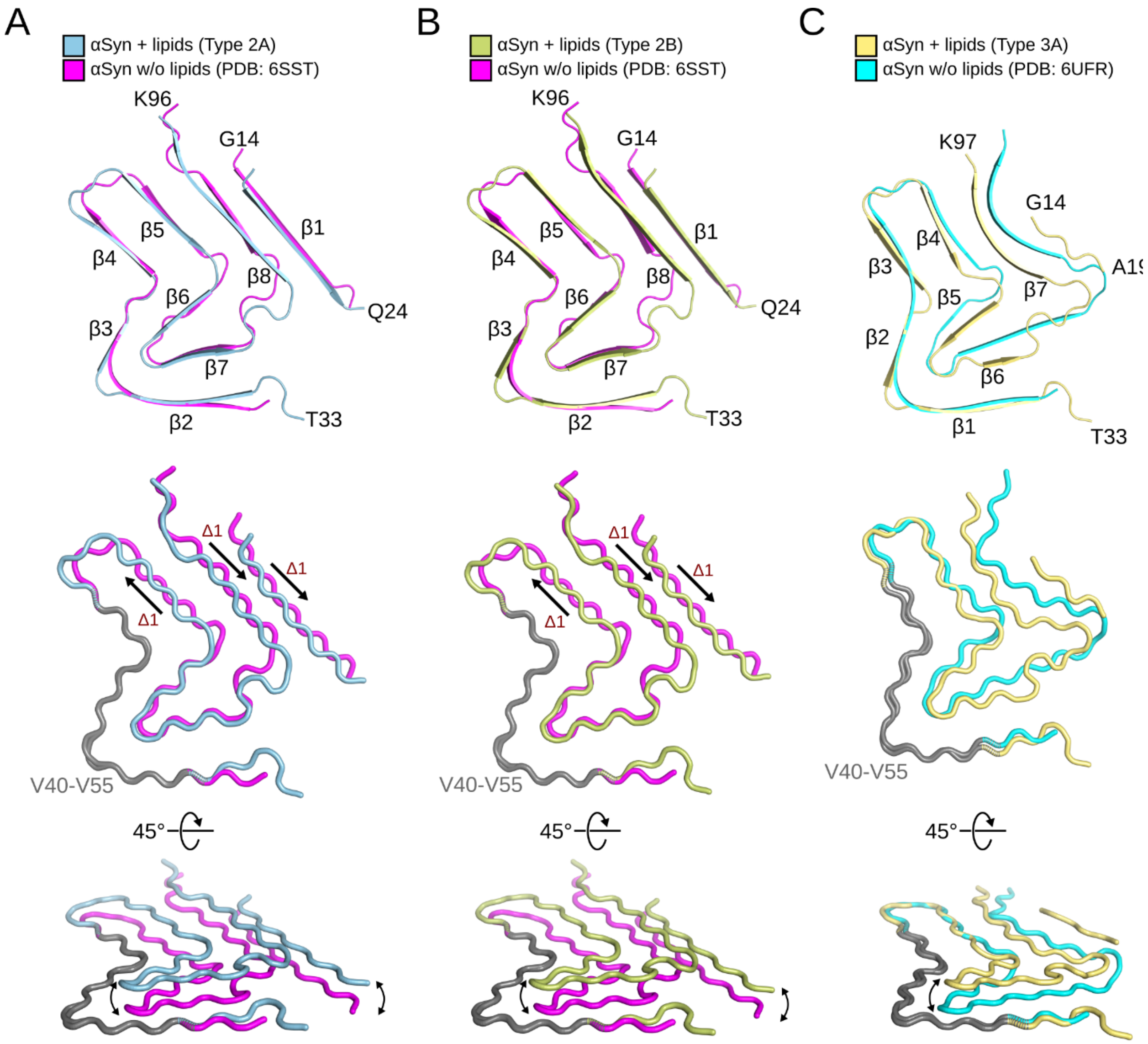


**Fig. S11. Comparison with between *L2* and *L3* fibrils and known structures.**
Superposition of a single protein chain of (**A**) *L2A* αSyn onto in vitro aggregated wild type αSyn (PDB: 6SST (*39*); Cα RMSD = 2.9 Å), (**B**) *L2B* αSyn onto in vitro aggregated wild type αSyn (PDB: 6SST (*39*); Cα RMSD = 3.0 Å), and (**C**) *L3A* αSyn onto in vitro aggregated E46K αSyn (PDB: 6UFR (*40*); Cα RMSD = 3.0 Å). Termini and β-strands are labeled. The middle panel visualizes the relative shift of β1, β5, and β8 introduced by the presence of lipids, after superimposing V40 to V55 (gray). The lower panel visualizes the out-of-plane shift of a single chain induced by the presence of lipids.

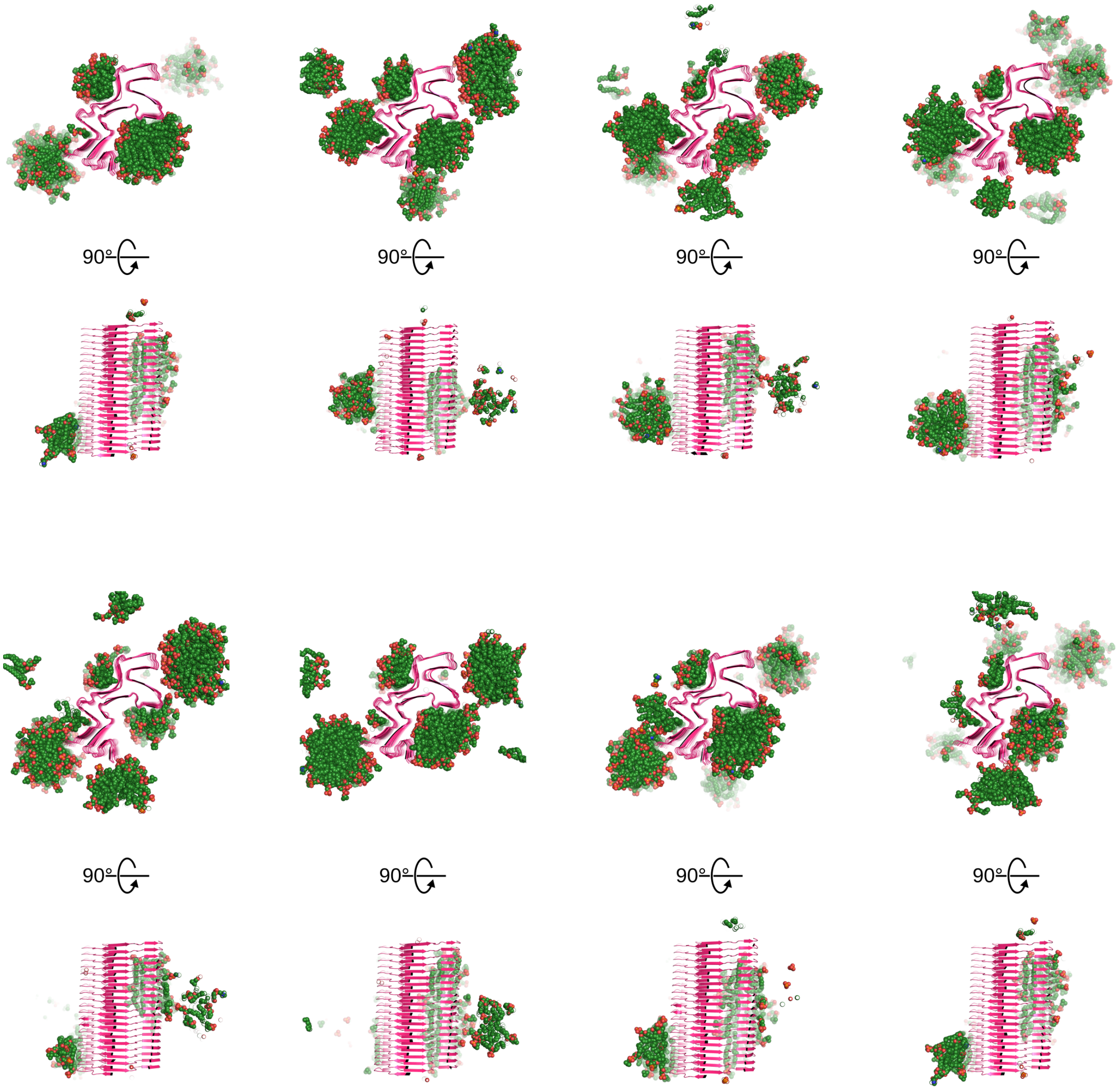


**Fig. S12. Molecular dynamics simulations of the lipidic *L1A* fibril.**
Cross-section through the conformations after eight 1 μs MD simulations of free-lipid diffusion in the presence of the lipidic *L1A* fibril, viewed from two perspectives. The fibril is shown as cartoon, the lipids as green spheres.

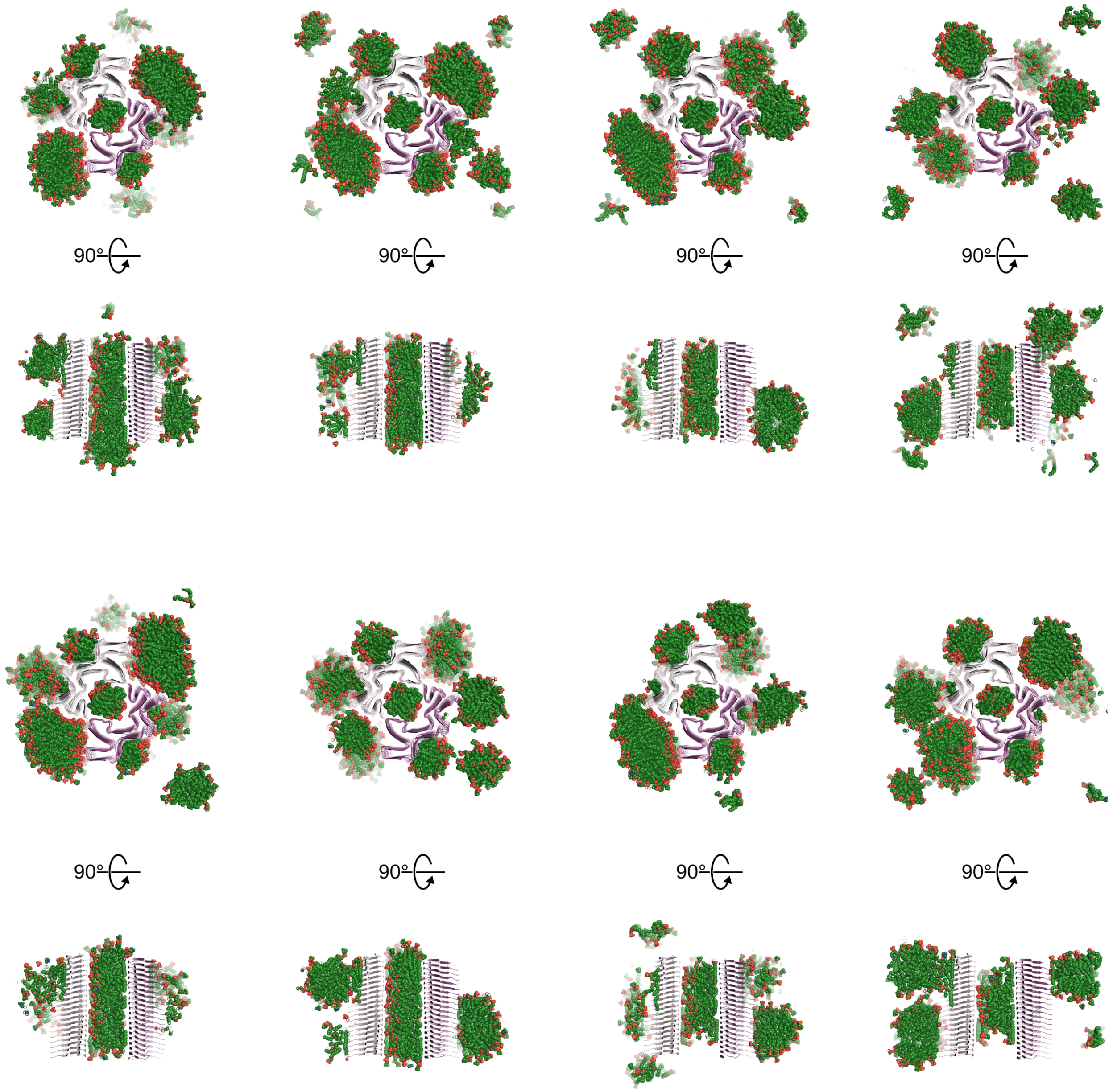


**Fig. S13. Molecular dynamics simulations of the lipidic *L1B* fibril.**
Cross-section through the conformations after eight 1 μs MD simulations of free-lipid diffusion in the presence of the lipidic *L1B* fibril, viewed from two perspectives. The fibril is shown as cartoon, the lipids as green spheres.

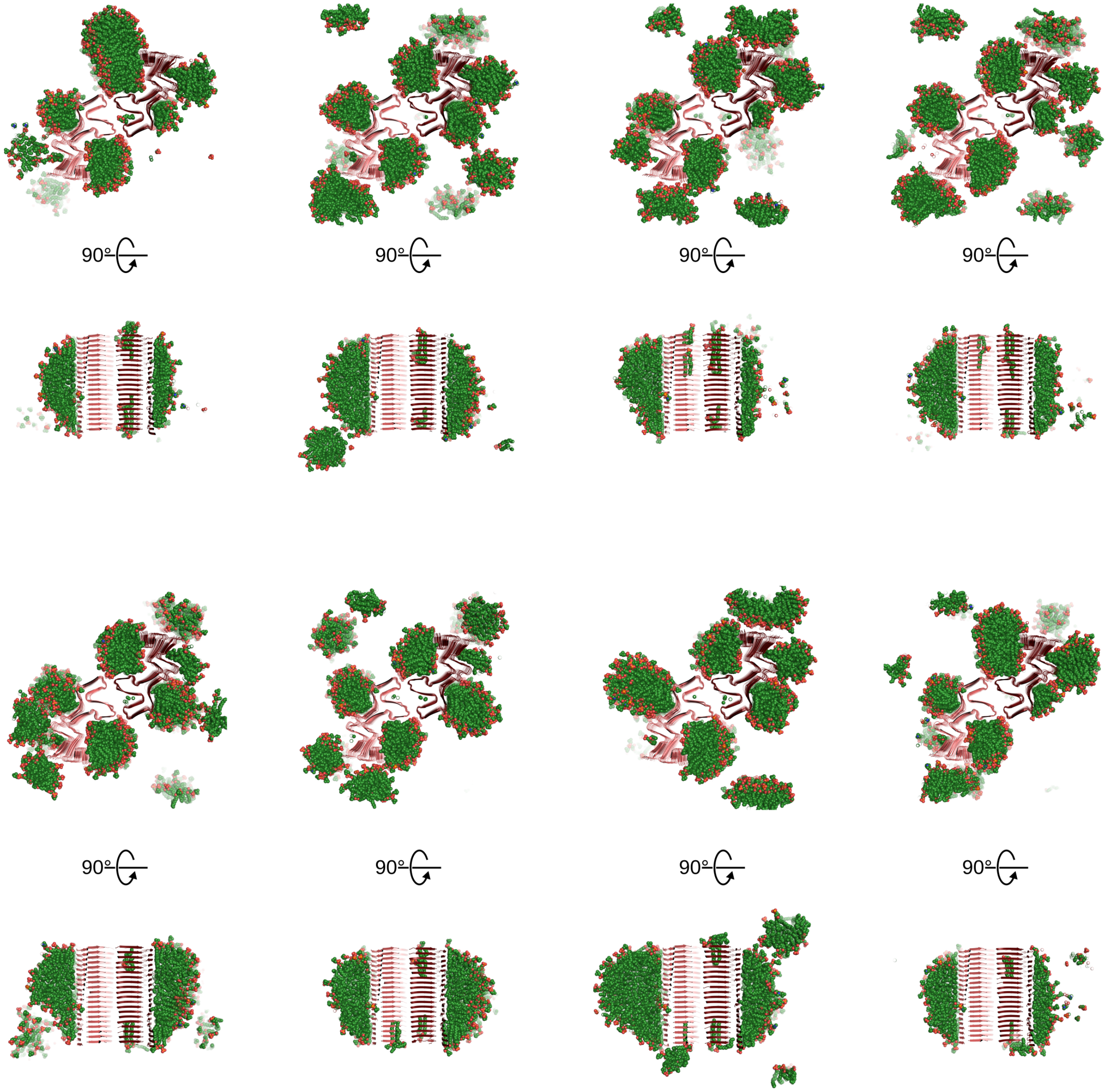


**Fig. S14. Molecular dynamics simulations of the lipidic *L1C* fibril.**
Cross-section through the conformations after eight 1 μs MD simulations of free-lipid diffusion in the presence of the lipidic *L1C* fibril, viewed from two perspectives. The fibril is shown as cartoon, the lipids as green spheres.

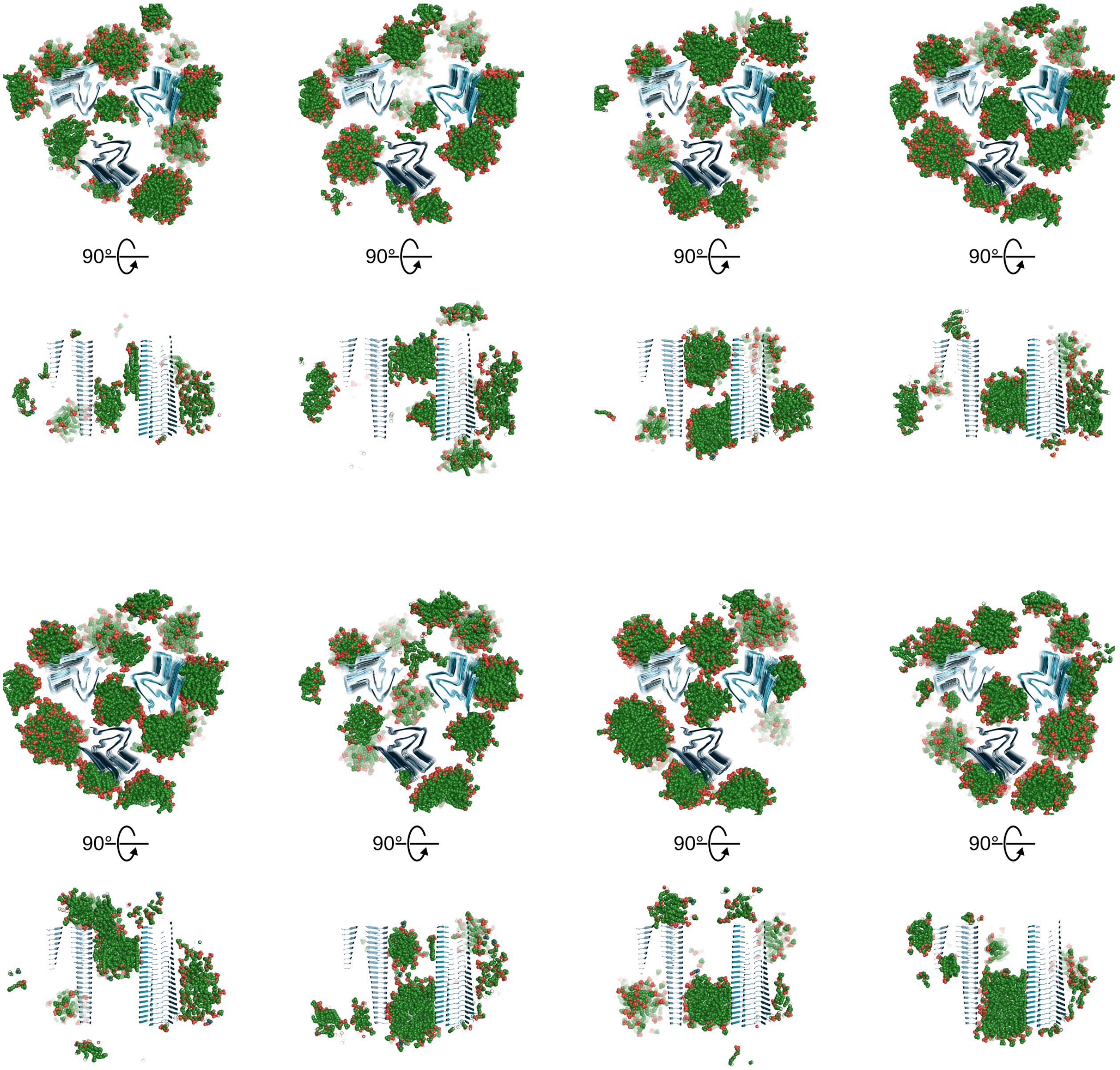


**Fig. S15. Molecular dynamics simulations of the lipidic *L2A* fibril.**
Cross-section through the conformations after eight 1 μs MD simulations of free-lipid diffusion in the presence of the lipidic *L2A* fibril, viewed from two perspectives. The fibril is shown as cartoon, the lipids as green spheres.

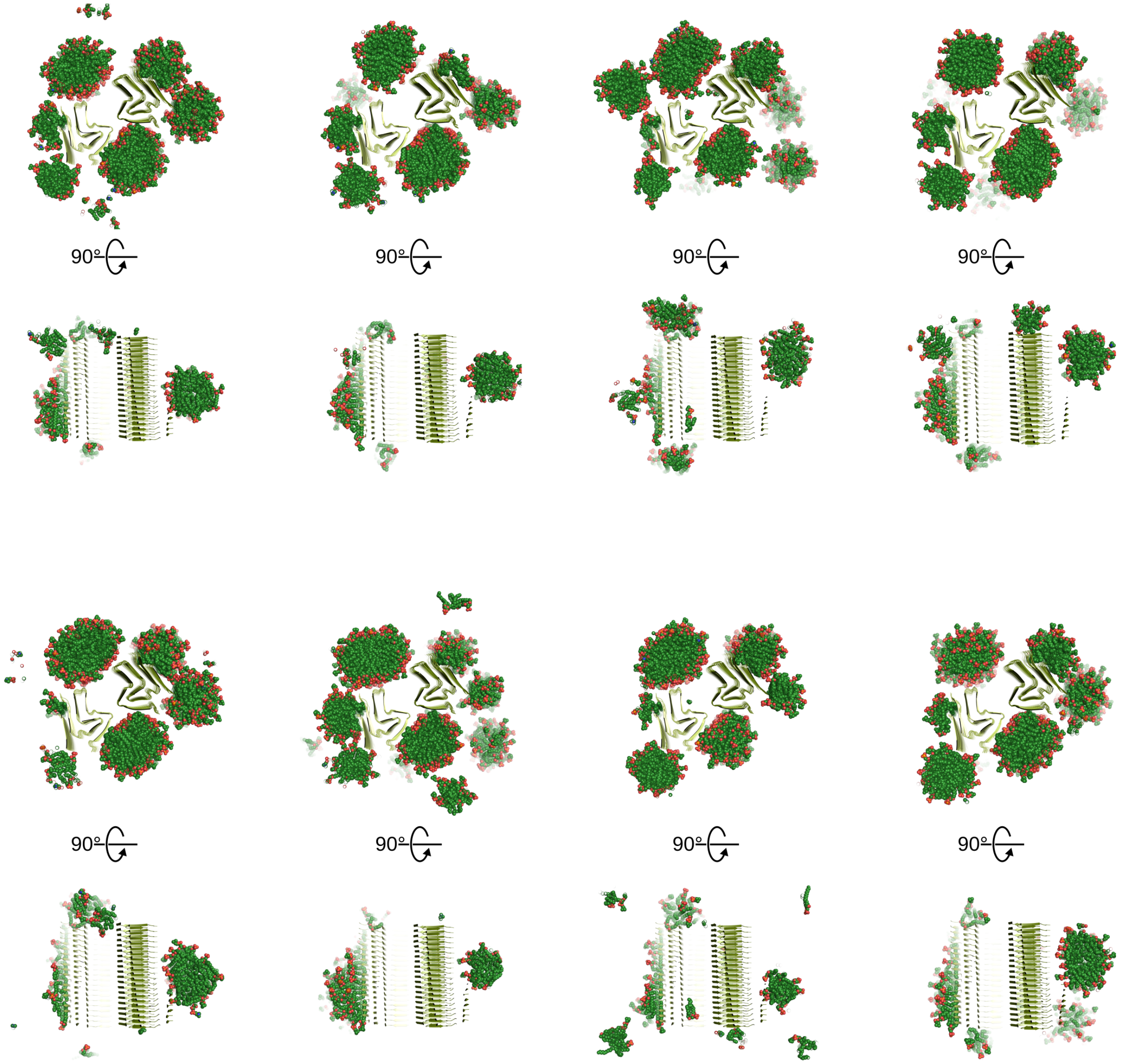


**Fig. S16. Molecular dynamics simulations of the lipidic *L2B* fibril.**
Cross-section through the conformations after eight 1 µs MD simulations of free-lipid diffusion in the presence of the lipidic *L2B* fibril, viewed from two perspectives. The fibril is shown as cartoon, the lipids as green spheres.

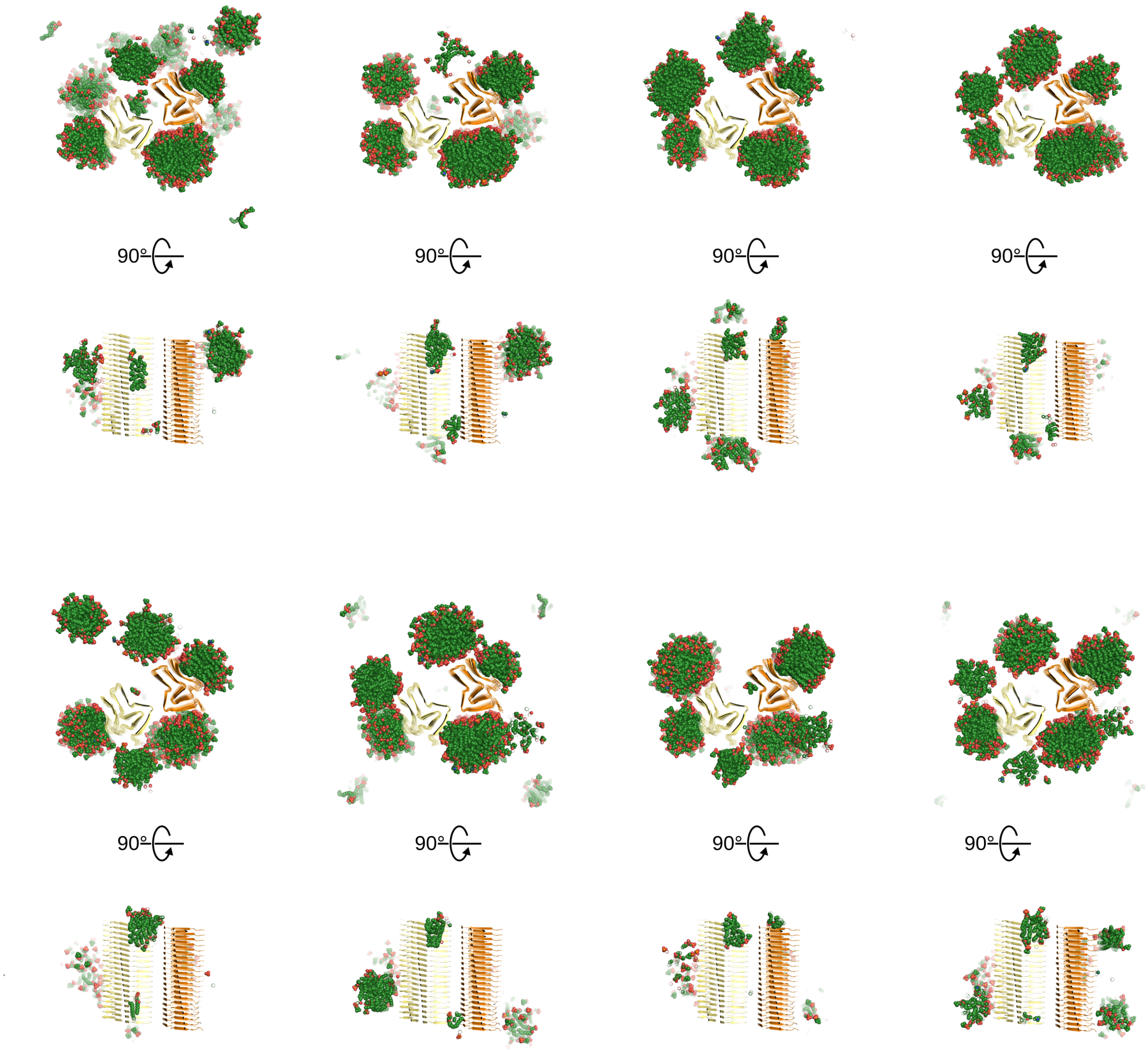


**Fig. S17. Molecular dynamics simulations of the lipidic *L3A* fibril.**
Cross-section through the conformations after eight 1 μs MD simulations of free-lipid diffusion in the presence of the lipidic *L3A* fibril, viewed from two perspectives. The fibril is shown as cartoon, the lipids as green spheres.

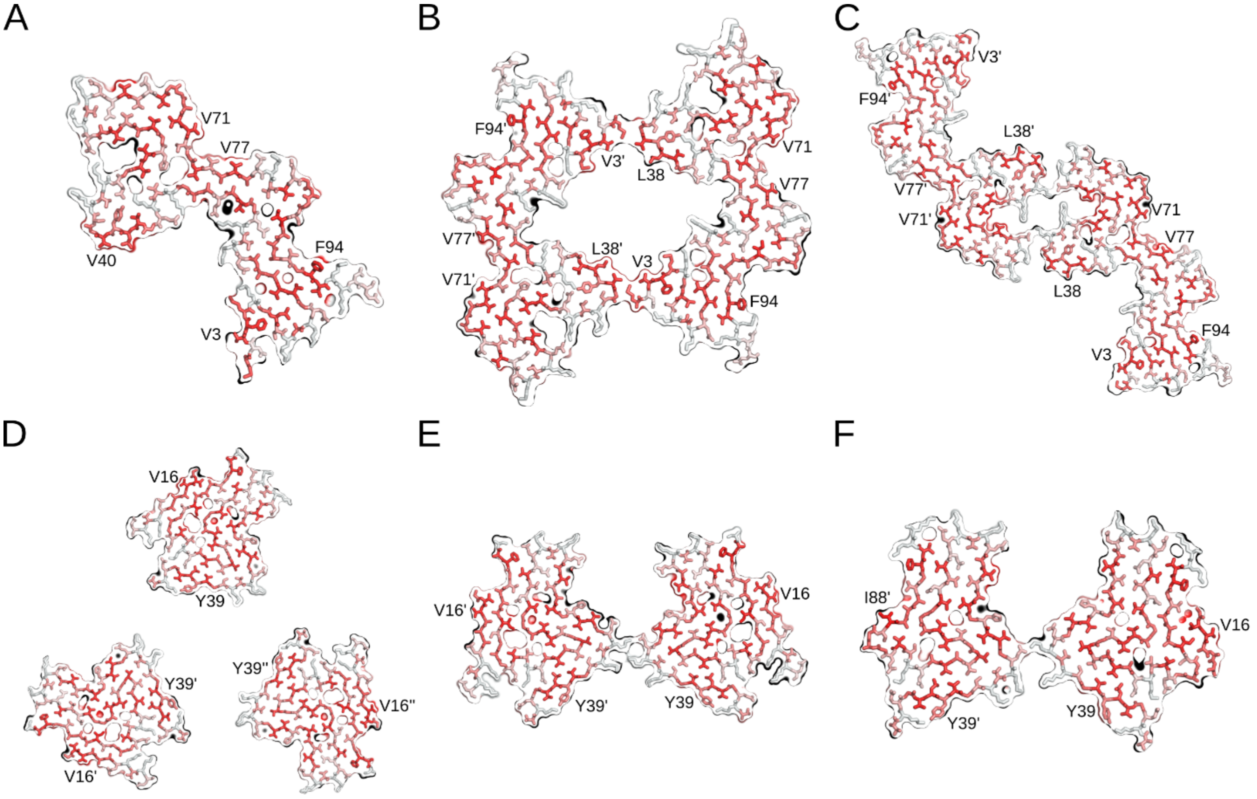


**Fig. S18. The hydrophobicity of lipid-induced αSyn fibrils.**

The atomic models of *L1A* (**A**), *L1B* (**B**), *L1C* (**C**), *L2A* (**D**), *L2B* (**E**), and *L3A* (**F**) αSyn fibrils with amino acids colored according to the Eisenberg hydrophobicity scale (*57*). Regions in red are hydrophobic. For orientation, surface amino acids in the center of a hydrophobic region are labeled.

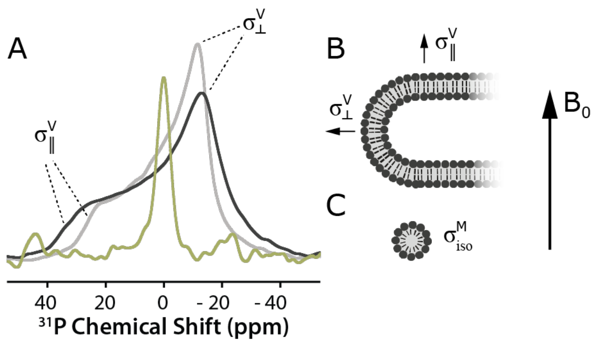


**Fig. S19. 31P ssNMR experiments for the determination of lipid aggregation state**

**A**: $^{1}H$ decoupled static $^{31}P$ ssNMR spectra of vesicles of POPA and POPC (1:1) at 280 K (black) and 310 K (grey) compared to the spectrum of lipidic αSyn fibrils at 280 K (green, same sample as dataset 2w). Spectra of vesicles show a characteristic powder pattern due to chemical shift anisotropy (CSA) after uniaxial diffusion of the lipid molecule about its own long axis. Lateral diffusion of lipid molecules does not result in significant reorientation, consistent with lipid bilayer structures of low curvature (**B**). Lipids bound to αSyn fibrils show a single sharp line, indicating that CSA is averaged via isotropic reorientation of lipid headgroup moieties, consistent with the presence of high-curvature lipid aggregates, such as micelles (**C**). This behavior cannot be explained by a change of the lipid transition temperature and a resulting increase in mobility, since even at higher temperatures the vesicle spectra do not show a comparably sharp line.

**Tab. S1. Cryo-EM structure determination statistics.**

| **Lipid-induced PM** | ***L1A*** | ***L1B*** | ***L1C*** | ***L2A*** | ***L2B*** | ***L3A*** |
|---|---|---|---|---|---|---|
| **Data collection** | | | | | | |
| Microscope | Titan Krios G2 | Titan Krios G2 | | | Titan Krios G2 | |
| Voltage [keV] | 300 | 300 | | | 300 | |
| Detector | K3 | K3 | | | K3 | |
| Magnification | 81,000 | 81,000 | | | 81,000 | |
| Pixel size [Å] | 1.05 | 1.05 | | | 1.05 | |
| Defocus range [µm] | -0.5 to -2.0 | -0.7 to -2.0 | | | -0.5 to -2.0 | |
| Exposure time [s/frame] | 2.997 | 2.997 | | | 2.997 | |
| Number of frames | 40 | 50 | | | 40 | |
| Total dose [$e^-$/Å$^2$] | 42.72 (1.07 $e^-$/Å$^2$/frame) | 50.83 (1.02 $e^-$/Å$^2$/frame) | | | 42.72 (1.07 $e^-$/Å$^2$/frame) | |
| **Reconstruction** | | | | | | |
| Micrographs | 4,589 | 4,324 | | | 4,542 | |
| Box width [pixels] | 250 | 250 | | | 250 | |
| Inter-box distance [pixels] | 14 | 14 | | | 14 | |
| Picked segments (no.) | 585,342 | 504,236 | | | 1,223,706 | |
| | *L1A* | *L1B* | *L1C* | *L2A* | *L2B* | *L3A* |
| PDB-ID | XXX | XXX | XXX | XXX | XXX | XXX |
| EMDB-ID | XXX | XXX | XXX | XXX | XXX | XXX |
| Final segments [no.] | 13,641 | 19,108 | 25,817 | 46,003 | 20,388 | 46,882 |
| Final resolution [Å] (FSC=0.143) | 3.24 | 2.98 | 2.95 | 2.68 | 3.05 | 2.76 |
| Applied map sharpening B-factor [Å$^2$] | -85.24 | -83.67 | -87.28 | -98.95 | -78.72 | -85.99 |
| Symmetry imposed | C1 | C1 | C2 | C3 | C1 | C1 |
| Helical rise [Å] | 4.69 | 2.37 | 4.69 | 4.68 | 4.69 | 4.72 |
| Helical twist [°] | -0.95 | 179.49 | -0.72 | -0.75 | -0.82 | -0.95 |

**Tab. S2. Model building statistics.**

| Lipid-induced PM | *L1A* | *L1B* | *L1C* | *L2A* | *L2B* | *L3A* |
|---|---|---|---|---|---|---|
| **Initial model [PDB code]** | *de novo* | *de novo* | *de novo* | 6SST | 6SST | 6SST |
| **Model composition** | | | | | | |
| Chains | 5 | 10 | 10 | 15 | 10 | 10 |
| Non-hydrogen atoms | 3,460 | 6,920 | 6,920 | 7,755 | 5,170 | 4,665 |
| Protein residues | 495 | 990 | 990 | 1,125 | 750 | 680 |
| **RMS deviations** | | | | | | |
| Bond lengths [Å] | < 0.01 | < 0.01 | 0.01 | < 0.01 | < 0.01 | 0.01 |
| Bond angles [°] | 0.82 | 0.64 | 1.5 | 0.65 | 0.42 | 1.16 |
| **Validation** | | | | | | |
| MolProbity score | 2.39 | 2.37 | 2.95 | 1.53 | 1.32 | 2.49 |
| Clashscore | 20.22 | 16.36 | 8.01 | 10.11 | 5.84 | 12.49 |
| **Ramachandran plot** | | | | | | |
| Outliers [%] | 0 | 0 | 0 | 0 | 0 | 0 |
| Allowed [%] | 11.34 | 9.28 | 7.73 | 0 | 0 | 8.46 |
| Favored [%] | 88.66 | 90.72 | 92.27 | 100 | 100 | 91.54 |

**Movie S1. Lipid binding to the *L1B* αSyn fibril.**

The movie shows the first 100 ns of a representative trajectory of randomly placed phospholipids (1:1 mixture of POPC/POPA) binding to the *L1B* αSyn fibril. The lipids are shown as green-sphere model, and the αSyn fibril as cartoon, with both protofilaments colored differently.

### Supplemental References


29. W. Hoyer *et al.*, Dependence of alpha-synuclein aggregate morphology on solution conditions. *J. Mol. Biol.* **322**, 383-393 (2002).
30. L. Antonschmidt *et al.*, Insights into the molecular mechanism of amyloid filament formation: Segmental folding of alpha-synuclein on lipid membranes. *Sci. Adv.* **7**, eabg2174 (2021).
31. A. Böckmann *et al.*, Characterization of different water pools in solid-state NMR protein samples. *J. Biomol. NMR* **45**, 319 (2009).
32. E. Barbet-Massin *et al.*, Rapid Proton-Detected NMR Assignment for Proteins with Fast Magic Angle Spinning. *J. Am. Chem. Soc.* **136**, 12489-12497 (2014).
33. D. N. Mastronarde, Automated electron microscope tomography using robust prediction of specimen movements. *J. Struct. Biol.* **152**, 36-51 (2005).
34. D. Tegunov, P. Cramer, Real-time cryo-electron microscopy data preprocessing with Warp. *Nat Meth* **16**, 1146-+ (2019).
35. J. Zivanov, T. Nakane, S. H. W. Scheres, Estimation of high-order aberrations and anisotropic magnification from cryo-EM data sets in RELION-3.1. *Iucrj* **7**, 253-267 (2020).
36. S. He, S. H. W. Scheres, Helical reconstruction in RELION. *J. Struct. Biol.* **198**, 163-176 (2017).
37. A. Rohou, N. Grigorieff, CTFFIND4: Fast and accurate defocus estimation from electron micrographs. *J. Struct. Biol.* **192**, 216-221 (2015).
38. P. Emsley, K. Cowtan, Coot: model-building tools for molecular graphics. *Acta Crystallogr D* **60**, 2126-2132 (2004).
39. R. Guerrero-Ferreira *et al.*, Two new polymorphic structures of human full-length alpha-synuclein fibrils solved by cryo-electron microscopy. *Elife* **8**, (2019).
40. D. R. Boyer *et al.*, The alpha-synuclein hereditary mutation E46K unlocks a more stable, pathogenic fibril structure. *Proc. Natl. Acad. Sci. U. S. A.* **117**, 3592-3602 (2020).
41. P. V. Afonine *et al.*, Real-space refinement in PHENIX for cryo-EM and crystallography. *Acta Crystallogr D* **74**, 531-544 (2018).
42. D. Liebschner *et al.*, Macromolecular structure determination using X-rays, neutrons and electrons: recent developments in Phenix. *Acta Crystallogr D* **75**, 861-877 (2019).
43. V. B. Chen *et al.*, MolProbity: all-atom structure validation for macromolecular crystallography. *Acta Crystallogr D* **66**, 12-21 (2010).
44. L. Martinez, R. Andrade, E. G. Birgin, J. M. Martinez, PACKMOL: a package for building initial configurations for molecular dynamics simulations. *J. Comput. Chem.* **30**, 2157-2164 (2009).
45. C. Tian *et al.*, ff19SB: amino-acid-specific protein backbone parameters trained against quantum mechanics energy surfaces in solution. *J. Chem. Theory Comput.* **16**, 528-552 (2020).
46. I. R. Gould, S. A.A., C. J. Dickson, B. D. Madej, R. C. Walker, Lipid17: A comprehensive AMBER force field for the simulation of zwitterionic and anionic lipids. *in press*, (2018).
47. I. S. Joung, T. E. Cheatham III, Determination of alkali and halide monovalent ion parameters for use in explicitly solvated biomolecular simulations. *J. Phys. Chem. B* **112**, 9020-9041 (2008).

48. S. Izadi, R. Anandakrishnan, A. V. Onufriev, Building water models: a different approach. *J Phys Chem Lett* **5**, 3863-3871 (2014).
49. B. Frieg *et al.*, Molecular mechanisms of glutamine synthetase mutations that lead to clinically relevant pathologies. *PLoS Comput. Biol.* **12**, e1004693 (2016).
50. B. Frieg, L. Gremer, H. Heise, D. Willbold, H. Gohlke, Binding modes of thioflavin T and Congo red to the fibril structure of amyloid-beta(1-42). *Chem. Commun.* **56**, 7589-7592 (2020).
51. C. W. Hopkins, S. Le Grand, R. C. Walker, A. E. Roitberg, Long-time-step molecular dynamics through hydrogen mass repartitioning. *J. Chem. Theory Comput.* **11**, 1864-1874 (2015).
52. J. P. Ryckaert, G. Ciccotti, H. J. C. Berendsen, Numerical integration of cartesian equations of motion of a system with constraints molecular dynamics of n-alkanes. *J. Comput. Phys.* **23**, 327-341 (1977).
53. T. Darden, D. M. York, L. G. Pedersen, Particle Mesh Ewald: an N·log (N) method for Ewald sums in large systems. *J. Chem. Phys.* **98**, 10089-10092 (1993).
54. D. A. Case *et al.*, AMBER 21. *University of California, San Francisco.*, (2021).
55. R. Salomon-Ferrer, A. W. Götz, D. Poole, S. Le Grand, R. C. Walker, Routine microsecond molecular dynamics simulations with Amber on GPUs. 2. Explicit solvent particle mesh Ewald. *J. Chem. Theory Comput.* **9**, 3878-3888 (2013).
56. D. R. Roe, T. E. Cheatham III, PTRAJ and CPPTRAJ: software for processing and analysis of molecular dynamics trajectory data. *J. Chem. Theory Comput.* **9**, 3084-3095 (2013).
57. D. Eisenberg, E. Schwarz, M. Komaromy, R. Wall, Analysis of membrane and surface protein sequences with the hydrophobic moment plot. *J. Mol. Biol.* **179**, 125-142 (1984).
